# GhostParser: A highly scalable phylogenomic approach for the identification of ghost introgression

**DOI:** 10.1101/2025.08.21.671585

**Authors:** Ethan R. Tolman, Anton Suvorov

## Abstract

A growing body of empirical research shows that interspecific gene flow is a widespread biological force that shapes evolutionary histories across the Tree of Life. Computational approaches designed to detect introgression either employ full likelihood, including Bayesian, frameworks to directly estimate phylogenetic networks or utilize summary statistics derived directly from locus sequence alignments or estimated gene trees to map gene flow events onto the species tree. Many current methods currently have major shortcomings. The computationally scalable summary statistics and pseudo-likelihood-based techniques may provide erroneous results in the presence of so-called “ghost” introgression and rate variation between lineages. On the other hand, full likelihood methods are more accurate, but are not computationally tractable for large phylogenomic datasets. Here, we develop a novel summary statistic, based on tree heights of different gene tree topologies to reliably distinguish between sampled and ghost introgression events. We implemented this approach in the publicly accessible bioinformatic pipeline “GhostParser”. We demonstrate that GhostParser can accurately distinguish between scenarios of sampled and ghost introgression, even in the presence of rate variation between lineages. Our methodology generally concurs in accuracy with the full likelihood software Bayesian Phylogenetics and Phylogeography (BPP) on empirical datasets, and outperforms BPP in our simulation conditions, both in a small fraction of the computational time. We show that GhostParser is a scalable tool for the identification of different introgression patterns in phylogenomic datasets.

## Introduction

A major shift in phylogenomics has been the realization that the evolutionary relationships oftentimes cannot be most accurately described by a binary tree model. Introgression is a source of novel genetic variation, which may be deleterious, adaptive or neutral and can cause the involved species or populations to share both genotypes and phenotypes (Whitney et al. 2006; Pardo-Diaz et al. 2012; Huerta-Sánchez et al. 2014; Bastide et al. 2018; Jones et al. 2018; Taylor and Larson 2019; Gibson et al. 2021; Hibbins and Hahn 2021). In a multi-locus dataset, introgression will produce an excess of loci supporting the same discordant history—that is, a higher proportion of gene trees will share an alternative topology that does not match the species tree. Such a signature of introgression has been identified in many lineages across the Tree of Life including vascular plants (Stull et al. 2023), mammals (Hallström and Janke 2010; Kumar et al. 2017; Teixeira and Cooper 2019; Gokcumen 2020; McArthur et al. 2021; Koller et al. 2022), birds (Singhal et al. 2021), fishes (Bernal et al. 2017; Qian et al. 2023), and arthropods (Leduc-Robert and Maddison 2018; Suvorov et al. 2022A,B; Thawornwattana et al. 2023). Indeed, as many as 10% of extant fauna and 25% of extant flora are estimated to be involved in introgression with other species (Mallet 2005). In each of these systems, and in all other clades where introgression has occurred, evolutionary history is better described as a phylogenetic network, than as a strictly bifurcating phylogenetic tree (Huson and Bryant 2006).

While many classical studies of introgression employ a “sampled introgression” model whereby gene flow is assumed to have involved two sampled lineages (Green et al. 2010; Edelman et al. 2019; Suvorov, et al. 2022A,B), this model does not fit every species network (Tricou, et al. 2022A,B). In many systems a “ghost introgression” model in which gene flow occurs from an unsampled lineage to a sampled lineage is more appropriate (Zhang et al. 2019; Ottenburghs 2020a; Chang et al. 2023; Pang and Zhang 2024; Zhang et al. 2024). Both models of introgression will result in an excess of loci supporting an alternative evolutionary history than a recovered species tree, and can also pose challenges for species tree estimation (Tricou et al., 2022A,B, Pang and Zhang 2023,2024). Distinguishing between models of ghost and sampled introgression is an important consideration in evolutionary biology, as an incorrectly applied introgression model can lead to erroneous conclusions regarding adaptation, speciation, and even the morphology of extinct lineages (Ottenburghs 2020b).

### Identifying introgression

The inference of introgression from phylogenomic datasets can be compromised by various factors. One of such factors is the among-lineage rate variation that increases branch length heterogeneity obscuring the signal of actual introgression in tests that utilize branch length, and creating a false positive signal of introgression between lineages in a multi-species network by making slower evolving lineages appear to have introgressed (Frankel and Ané 2023; Koppetsch et al. 2024).

The vast majority of current computational approaches designed to estimate reticulate evolution either utilize summary statistics derived from multiple sequence alignments (MSA) or gene trees to map events onto an estimated species tree (Edelman et al. 2019; Hahn and Hibbins 2019; Malinsky et al. 2021; Suvorov,, et al. 2022A), directly infer a phylogenetic network in either a pseudo-likelihood (Solís-Lemus et al. 2017; Allman et al. 2019) or in full-likelihood (including Bayesian) frameworks (Hey et al. 2018; Wen and Nakhleh 2018; Zhang et al. 2018; Flouri et al. 2020b Pang and Zhang 2024). These current methods offer a major tradeoff in accuracy and computational feasibility (Hibbins and Hahn 2022; Pang and Zhang 2024; Pfeifer et al. 2024).

#### Full likelihood approaches

Full likelihood methods that leverage multilocus sequence data directly, consider the joint distribution of coalescent times and gene-tree topologies to infer phylogenetic networks (Jiao et al. 2021). The full likelihood approach implemented in Bayesian Phylogenetics and Phylogeography (BPP) program (Flouri et al. 2020a) is considered to be the most accurate tool to identify introgression in multi-locus datasets. Furthermore, BPP provides the most accurate framework for distinguishing between ghost and sampled introgression models based upon testing in simulated and empirical datasets (Pang and Zhang 2024). BPP considers the joint probability distribution of gene-tree topologies and coalescent times under sampled and ghost introgression, using the foundational multi species coalescent with introgression (MSci) model (Flouri et al. 2020a). The marginal likelihoods of ghost, sampled inflow and outflow introgression models are compared for a given dataset to select the best fitting model, and identify gene flow involving sampled and ghost lineages. This technique has been demonstrated to be robust in the presence of between lineage-rate variation (Pang and Zhang 2024). However, even though BPP is potentially more scalable than other implementations of full likelihood approaches to identify introgression (Hey et al. 2018; Wen and Nakhleh 2018; Zhang et al. 2018; Flouri et al. 2020b; Pfeifer et al. 2024), computational resources remain a major barrier in its implementation. For instance, BPP, will require approximately 6 hours of real compute time to compare three different introgression scenarios for one species triplet with 1000 loci of 1000 basepairs each (Pang and Zhang 2024), making it unscalable for large taxon and/or locus sampling.

#### Pseudo-likelihood approaches for detecting introgression

Alternatively, pseudo-likelihood methodologies which estimate a phylogenetic network from gene trees are moderately less computationally intensive than the full likelihood approaches (Yu et al. 2014a). These methods estimate a phylogenetic network utilizing gene tree topologies within a (pseudo-)likelihood or distance-based framework (Yu et al. 2014a; Solís-Lemus et al. 2017; Pang and Zhang 2024). They all evaluate the pseudo-likelihood functions of different phylogenetic networks in order to identify the one that best explains the observed gene tree discordance under the multispecies network coalescent (MSNC) model. Implementations include InferNetwork_ML and InferNetwork_MPL from PhyloNet (Yu et al. 2014b; Yu and Nakhleh 2015), NANUQ from MSCquartets (Allman et al. 2019) and SNaQ from PhyloNetworks (Solís-Lemus and Ané 2016). Although, these pseudo-likelihood approaches are capable of differentiating between ghost and sampled introgression, they are sensitive to model misspecification (Pang and Zhang 2024) and can also falsely infer introgression between lineages when rate variation between lineages is present (Frankel and Ané 2023).

#### Heuristic approaches for detecting introgression

In comparison with full- and pseudo-likelihood approaches, heuristic based techniques for detecting introgression are highly computationally scalable, and can analyze hundreds of triplets or quartets with thousands of loci in a matter of minutes. Despite such appealing scalability these approaches are oftentimes less accurate and incapable of identifying ghost introgression. Typically, heuristic approaches rely on various summary statistics which can be obtained from either MSAs or gene trees. First, site pattern-based methods (e.g. the D-statistic (Green et al. 2010; Zheng and Janke 2018), HyDe (Blischak et al. 2018), and Dfoil (Pease and Hahn 2015)) compute certain asymmetries in pattern frequencies, that can be indicative of introgression. Second, gene tree topology count-based techniques (e.g. the Discordant Counts Test (Suvorov, et al. 2022A)) assess species tree-gene tree heterogeneity. Finally, branch length-based approaches (e.g. the Branch Length Test (Suvorov et al. 2022A), D3 (Hahn and Hibbins 2019), and QuiBL (Edelman et al. 2019)) utilize branch length distributional differences. Typically, the counts or branch lengths in the most abundant discordant topology are compared to those in the least abundant discordant topology. These methods can all flag possible gene flow patterns across an input species tree in a matter of minutes to hours in a large multilocus dataset, but they cannot distinguish between ghost and sampled introgression accurately, if it all, and oftentimes erroneously identify ghost introgression as sampled (Tricou et al., 2022A,B; Pang and Zhang 2024). Heuristic tests can also exhibit elevated rates of false positives for sampled introgression in the presence of between-species rate variation, nevertheless topology count-based methods are less susceptible to producing false positives than methods, which rely on branch lengths (Frankel and Ané 2023; Koppetsch et al. 2024).

#### A new approach for distinguishing ghost and sampled introgression

Here, we developed a new heuristic-based strategy to detect sampled and ghost introgression that relies on a simple gene tree summary statistic based on the averaged tree height. Additionally, we provide theoretical justification for its usage and show how it reduces biases associated with heterogeneity of rate variation among lineages. Our approach can correctly identify ghost introgression in simulated datasets with high accuracy, specificity and sensitivity. This method concurs with BPP (the only introgression detection software which is currently *k*nown to be unaffected by rate variation and can accurately distinguish between sampled and ghost introgression, but is limited in its application by the computational time necessary to run it) in empirical datasets where full-likelihood and other summary-statistics based methods failed. Our approach also differentiates between various introgression models with higher accuracy than BPP in our simulated scenarios. We use our technique to revisit several hypothesized instances of introgression, and demonstrate that some of these “classical” reticulation events are likely instances of ghost introgression. Our approach is implemented in the bioinformatic software GhostParser, which will be a useful tool for evolutionary biologists to disentangle sampled and ghost introgression in a scalable manner.

## Materials and Methods

### Conceptual framework

Here, we provide theoretical foundations for our triplet-based statistical tests that will be used to estimate introgression under different multispecies coalescent models. The tests of introgression for our method primarily rely on the disparity of gene tree frequencies as wel as their expected heights, i.e. *T*_*MRCA*._ For all the models considered here we establish two fundamental assumptions: (i) evolutionary history between taxa is known in a form of a phylogenetic tree and (ii) only one gene copy is sampled from the extant taxa.

For the first null model (**Fig 1a**), which is a traditional multispecies coalescent (MSC) framework (Rannala and Yang 2003), we assume that there are three species, *A,B* and *C* that are related by the spe ies tree topology *S* ((*A,B*), *C*) Further, we specify divergence times 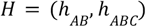 and let 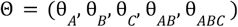 be the population-scaled mutation rate, i.e. θ 4*N*_*e*_ μ (Ewens 2004), where *N*_*e*_ and μ are effective population size and mutation rate, respectively. Both *H* and Θ are expressed n the expected number of mutations per site. Additionally, we define the probability (Rosenberg 2002) that two lineages (the number of lineages denoted as the superscript of *P*) do *not* coalesce (denoted as 0 in the subscript of *P*) in a population (e.g *AB*) as:

**Fig. 1.**
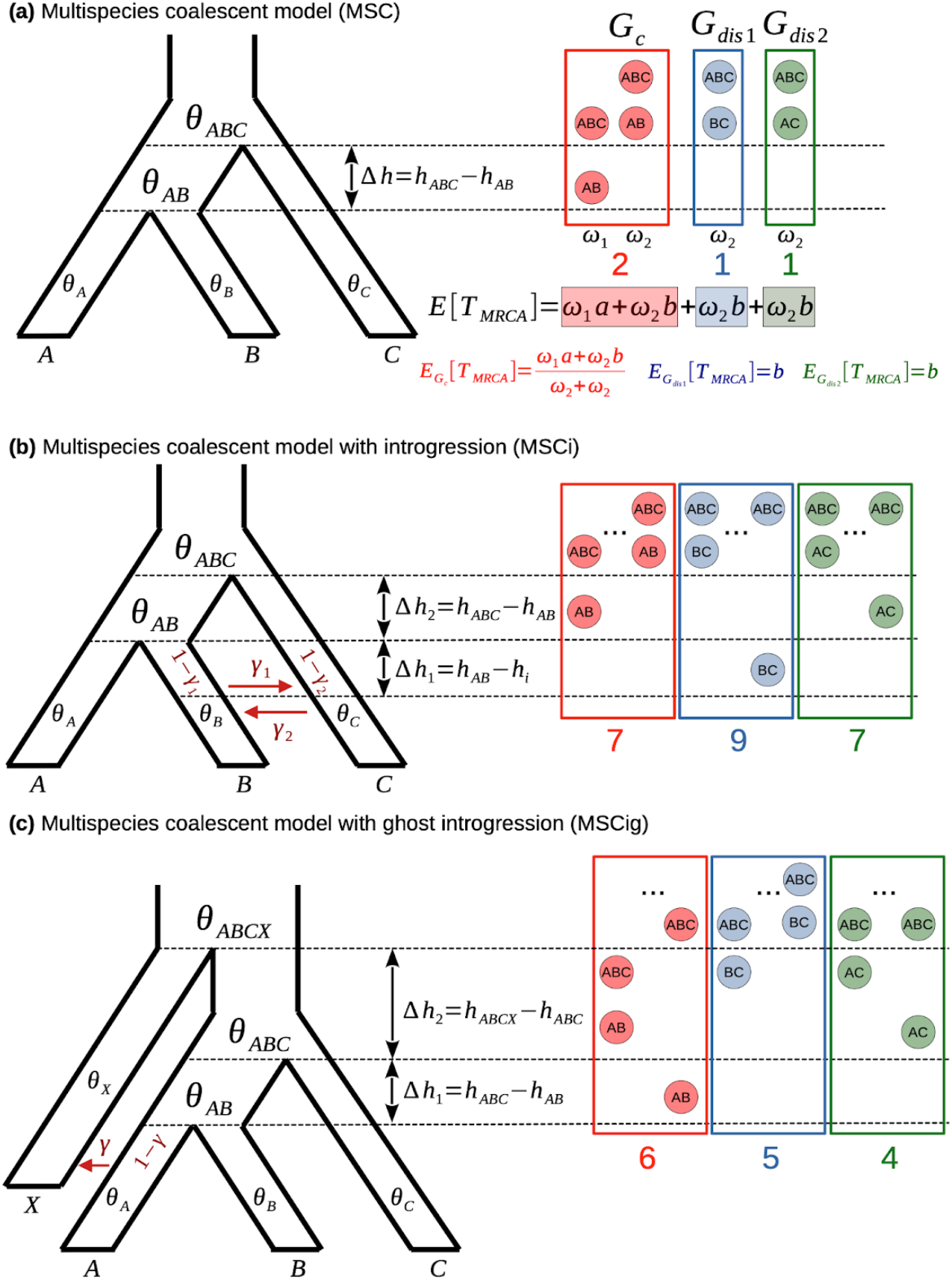
Parametrizations of GhostParser models: a) Multispecies coalescent (MSC) model of three species *A, B* and *C* with the corresponding population parameters 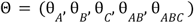 placed within the species tree topology ((*A B*) *C*).The arrow between dashed lines represents the duration Δ*h* of the branch calculated as the difference between divergence times *h*_*AB*_ and *h*_*ABC*._ The circles on the right denote the lineages (i.e. *AB* and *ABC*) and populations in which the first and second coalescence events of each triplet occur for concordant *G*_*c*_ ((*A,B*), *C*) as wel as discordant *G*_*dis1*_ ((*B,C*),*A*) and *G*_*dis2*_ ((*A,C*),*B*) topologies grouped by red, blue and green rectangles, respectively. The ω parameters represent probabilities at which such coalescence event configurations occur. The numbers below each rectangle show the number of ω parameters for concordant and two discordant topologies (see Appendix for more details). 𝔼 [*T*_*MRCA*_] represents the expected gene tree height that an be viewed mixture weighted by ω parameters, where *a* and *b* denote tree height for a specific gene tree. 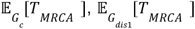,and 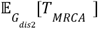 represent normalized expected values for gene tree heights of concordant and discordant topologies, respectively. **b)** Multispecies coalescent model with bidirectional introgression (MSCi) between non-sister species B and C. Red arrows indicate inflow and outflow introgression, with the introgression (“inheritance”) probabilities, γ_1_ and γ_2_ respectively. **c)** Multispecies coalescent model with ghost introgression between the extinct (unsampled) lineage *X* and extant species *A*. In this case only outflow introgression with probability γ_2_ is considered.

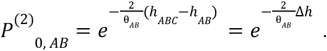

Thus, under the standard multispecies coalescent model (Pamilo and Nei 1988; Maddison 1997; Hudson 2002), the probability of concordant *G*_*c*_ = ((*A,B*), *C*) and discordant *G*_*dis1*_ =((*B, C*) *A*) and *G_*dis2*_* =((*A, C*), *B*) gene tree topologies are:

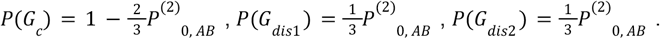

Obviously, under MSC the probabilities of concordant topology is larger than any of discordant topologies, i.e. *P*(*G*_*c*_) > *P*(*G* _*dis1*_) or *P*(*G*_*dis2*_) Additionally, the probabilities of discordant topologies are equal, i.e. *P*(*G* _*dis1*_) *P*(*G*_=*dis2*_). These probability comparisons for our null model provide a foundation for topology frequency tests (see below; Suvorov, et al. 2022A).

Now we consider the expected tree height *T*_*MRCA*_ for a gene tree in a single population with a constant *N*_*e*_ which can be expressed as 𝔼 [*T*_*MRCA*_]= 𝔼 [*T*_1_ + *T*_2_ + · · · + *T*_*n* − 1_ + *T*_*n*_] where *T* is the waiting time until next coalescence and its subscript denotes the total number of lineages *k* before a coalescence event that will reduce them to *k* − 1 (Wakeley 2009). The distribution of waiting time *t* until a coalescent event exhibits an exponential density:

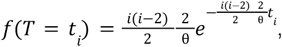

with the 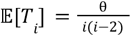 (Wa*k*eley 2009).

In MSC, however, in order to compute 𝔼 [*T*_*MRCA*_] we need to account for the probability ω of each gene tree and population parameters that can vary among species tree’s branches (see Appendix for more details). Thus, the expected gene tree height under the MSC model can be viewed as a weighted mixture. Define 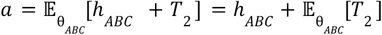 and 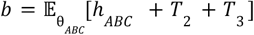, then:

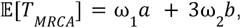

where each mixture component corresponds to the expected gene tree height multiplied by its specific probability ω, noting that 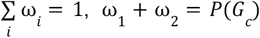 and 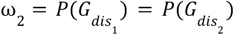. Now we evaluate all pairwise comparisons of 𝔼 [*T*_*MRCA*_] between discordant and concordant topologies, i.e. 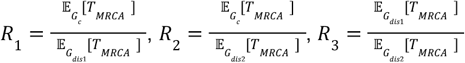.

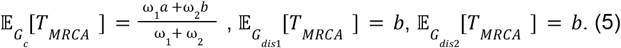

In this null case 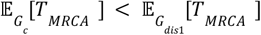 since *b* > *a* unless the probability of ω_1_= *P*^(2)^_0,AB_ ⟶0 i.e. for the internal branch length Δ*h* ⟶ 0 or θ_*AB*_ ⟶ ∞. The same result follows for the 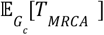 vs. 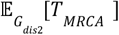 comparison. Additionally, under MSC the mixture components for discordant topologies *G*_*dis1*_ and *G*_*dis2*_ are expected to have equal expectations for gene tree heights. These disparities between 𝔼[*T*_*MRCA*_]’s for various gene tree topologies serve as foundation for the tree height test (THT) comparison (see below).

For the second scenario (**Fig 1b**), i.e. the multispecies coalescent with introgression model (MSCi) (Yu et al. 2014c), we allow a bidirectional gene flow to occur between non-sister species *B* and *C* with the probabilities of outflow and inflow Γ= (γ _1_,γ _2_) Specifically, for instance the γ _1_ can be interpreted as the probability of a gene copy in species *B* “slipping” via introgression “bridge” into the population of species *C* whereas 1 −γ_1_ will correspond to the probability of gene copy in species *B* staying in population *B* Note that under the formulation of the MSCi model here the introgression takes place only at fixed time points, thus it represents a special case of a more general multispecies network coalescent model (Zhu and Degnan 2017). As in MSC we define 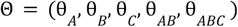 and 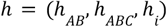 *h*) where *h*_*i*_ denotes not a divergence time but rather a time of the introgression event; hence, if gene copy *B* ends up in population *C* the probability that gene copies *B* and *C* don’t coalesce in population *C* (and vise versa) is:

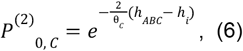

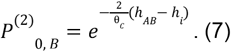

Following similar logic as for MSC, we can derive probabilities of concordant and discordant topologies:

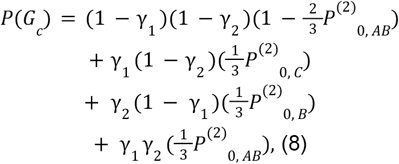

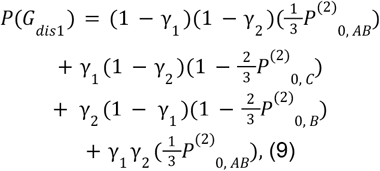

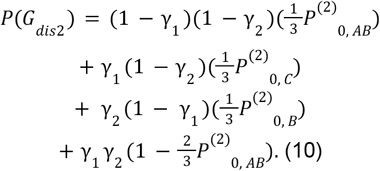

In contrast to the MSC, the probability of discordant topologies can differ, specifically:

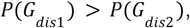

only if

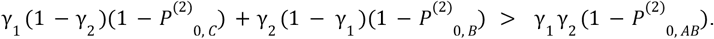

It is straightforward to show that for the special case of a unidirectional introgression i.e. γ_1_ = 0 (or γ_2_=0), the *P*(*G*_*dis1*_) > *P*(*G*_*dis2*_) will hold unless *P*^(2)^_0, *B*_ →1 (or *P*^(2)^_0,C_ ⟶ 1).

Now we define the 𝔼 [*T*_*MRCA*_] for the MSCi model (see Appendix for more details) and denote

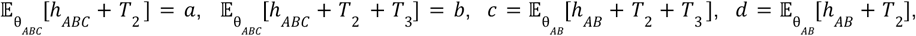

where *d* < *c* < *a* < *b*, then:

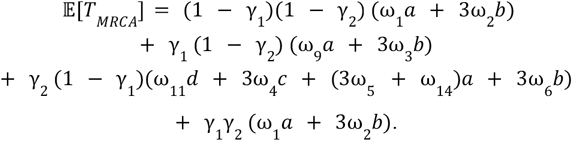

In the next step we examine all pairwise comparisons of weighted expectations like in the MSC case. Contrary to the MSC model, however, the *R*_1_ *R*_2_ and *R*_3_ expectation ratios derived from MSCi may exhibit non-trivial dependency on the combination of model parameters (**Fig. 2, supplementary Fig. S1**). Based on this observation, we need to determine which of those ratios has the highest power to detect introgression, i.e., the ratio that on average deviates from unity the most. To that end, we performed 10^5^ Monte Carlo simulations to evaluate to what extent each ratio deviates from one by calculating difference between expectation ratio and one, i.e. *R* − 1 For each iteration, to simulate a value of the expectation ratio we sampled a set of model parameters from uniform distributions *U*(0, 1) (**Fig. 3**). The simulation results indicate that *R*_1_ has the highest deviation from one (mean = 0.184 and median = 0.091). This suggests that comparisons of gene tree height expectations derived from concordant topologies 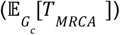 and discordant topologies 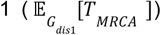 will li*k*ely exhibit higher statistical power (i.e. the probability of correctly rejecting the null MSC model if MSCi is true) compared to other expectation ratios. Additionally, we estimated how many times each expectation ratio is greater or smaller than one. Analyzing the simulation results we found that *R*_1_ > 1 in ∼90 9% *R*_2_ > 1 in ∼49 9% and *R*_3_ > 1 in ∼9%.

**Fig. 2.**
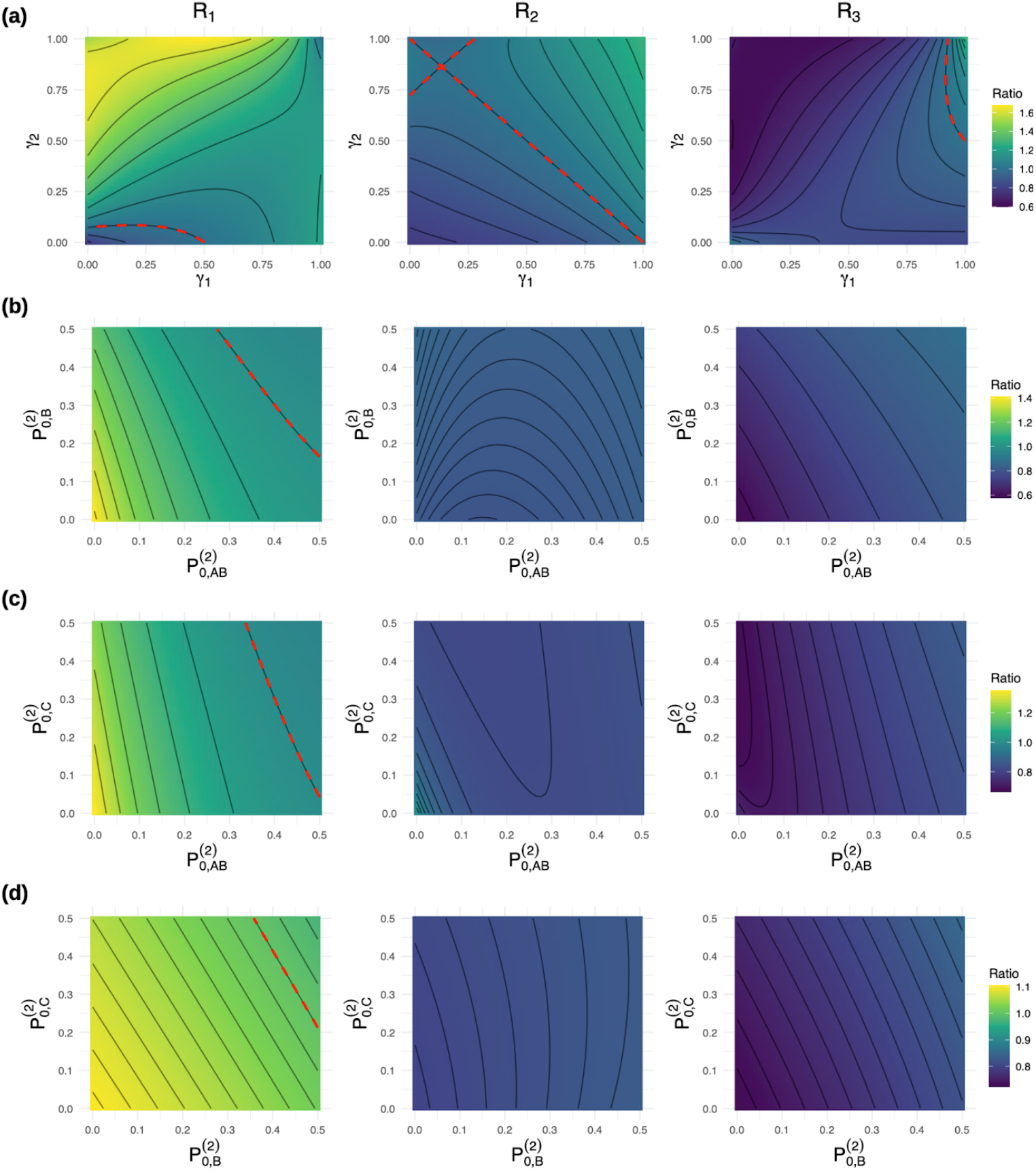
Contour plots of the expectation surface under MSCi model. a)- d) The changes of expectation ratio value in respect to the varying parameters of the MSCi model. The *R*_1_ trough *R*_3_ represent expectation ratios, i.e. 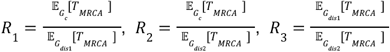.The red dashed line mar*k*s the region where expectation ratios attain unity.*P*^(2)^ _0,*AB*_ *P*^(2)^ _0, *B*_ and *P*^(2)^_0,*C*_ denote the probabilities of two lineages not coalescing in the populations *AB, B* or *C*, respectively. The γ_1_ and γ_2_ parameters represent introgression probabilities. Only two parameters were being varied at a time, while the rest were fixed at γ_1_=0 1 γ_2_=0.1, *P*^(2)^_0,*A*_= *P*^(2)^_0,B_ = *P*^(2)^_0,C_ =0.3, *a*= 0. 3, *b* =0.4, *c* =0.2, *d*= 0. 1.

**Fig. 3.**
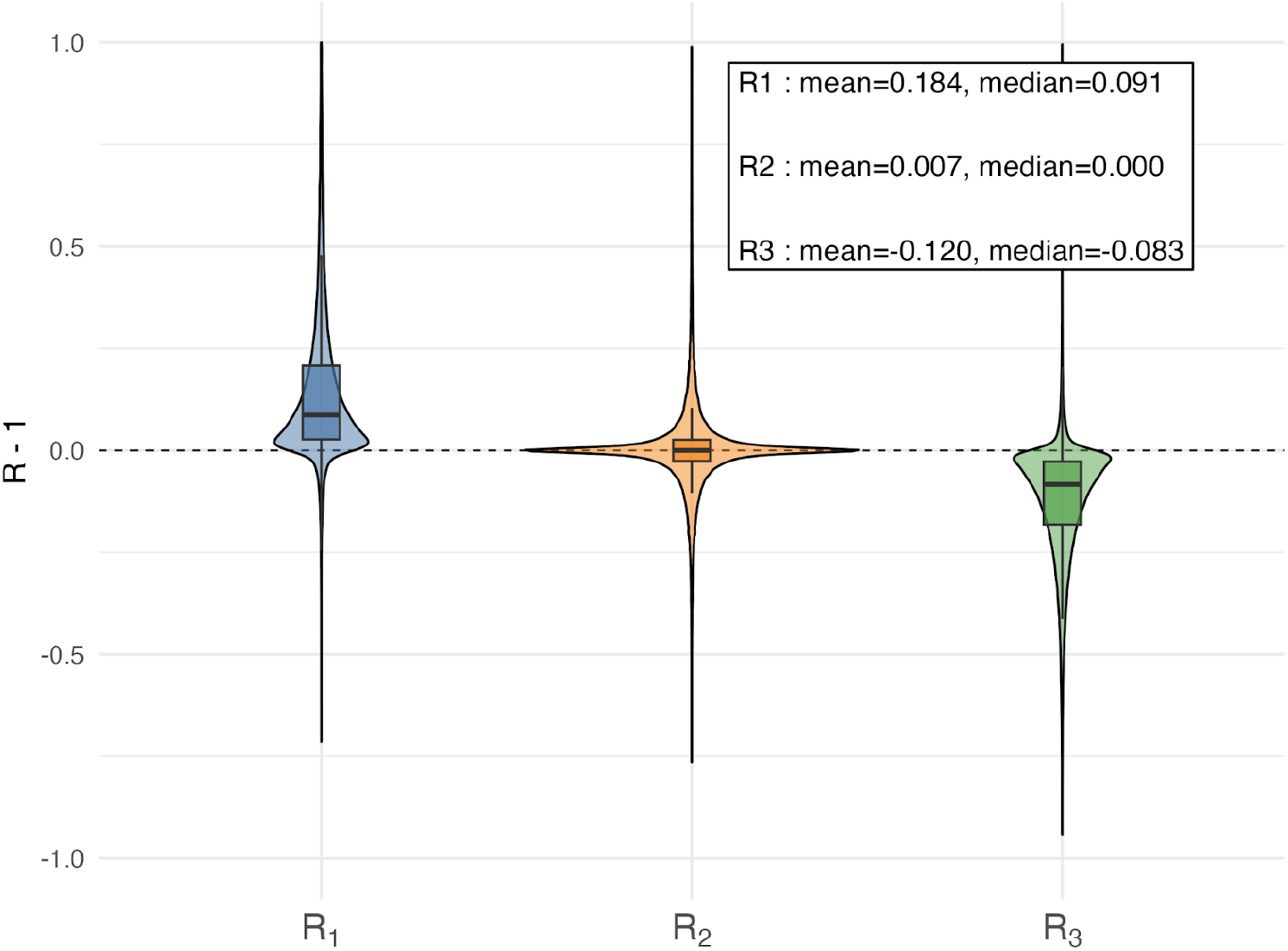
Distribution of simulated expectation ratios under MSCi: Violin plots represent simulated distributions of the *R*− 1 values for each of the expectation ratios, i.e. 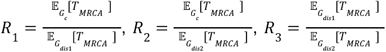. The box plot within each violin shows median as well as lower and upper quantiles. The dashed line marks the *R* − 1 values of zero.

Finally, we consider the multispecies coalescent with introgression scenario (MSCig) where gene flow occurs exclusively between the contemporary lineage *A* and an extinct/unsampled (“ghost”) lineage *X* (**Fig. 1c**). Furthermore, we assume that there is no introgression between neither *C* and *A* nor *C* and *B* Since the gene copy from *X* is not sampled, only outflow introgression from *A* to *X* is parametrized by probability γ For this model we define 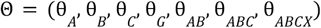 and 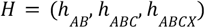 Given the parametrization of the MSCig we can derive frequencies of concordant and discordant gene tree topologies as follows:

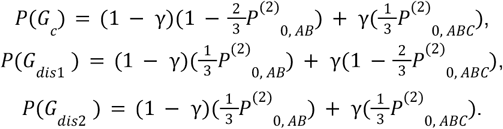

In the case of the MSCig model the difference of discordant topology probabilities will not depend γ (unless γ = 0 which will reduce MSCig to MSC null model where *P*(*G* _*dis1*_) = *P*(*G*_*dis2*_)) or

*P*_0, *AB*_, i.e.

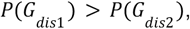

only if

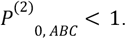

In other words, the frequency of discordant gene tree topologies *G*_*dis1*_ will be always greater than frequency of *G*_*dis1*_ unless θ_*ABC*_ → ∞ or Δ*h* = *h*_*ABCX*_ − *h*_*ABC*_ → 0.Next we formulate the 𝔼[*T* _*MRCA*_] for the MSCig model (see Appendix for more details) with 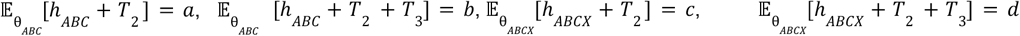 where *a* < *b* < *c* < *d*, then:

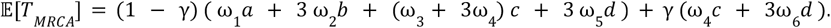

As in the case of the MSCi model the expectation ratios have complex relationships with the combinations of model parameters (**Fig. 4**). Again, based on the Monte Carlo simulation for the MSCig model, we determined that the *R*_2_ ratio only marginally outperformed *R*_1_ ratio using mean and median criteria of *R* − 1 distributions (**Fig. 5**). As for the MSCi model, here we calculated how many times the expectation ratios are greater or smaller than one in simulations. This analysis yielded *R* _1_> 1 in ∼14 9% *R*_2_ > 1 in ∼0 7% and *R* _3_> 1 in ∼29 2% Nevertheless, because the same gene tree height expectation comparisons must be used consistently across all three models (MSC, MSCi, and MSCig), we selected the *R*_1_ ratio as the basis. *R*_1_ shows markedly larger deviations under MSCi than *R*_2_ which in turn performs worse than *R*_3_ Moreover, because the values of *R*_1_ tend to exceed one under MSCi but fall below one under MSCig (Fisher’s exact test, odds ratio = 56.76, 95% CI: 55.2-58.3, *P* ∼ 0) in simulations, this property can be used to distinguish the two models: 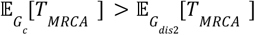 for MSCi, whereas 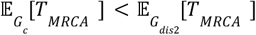 for MSCig with a success probability of ∼78.7%.

**Fig. 4.**
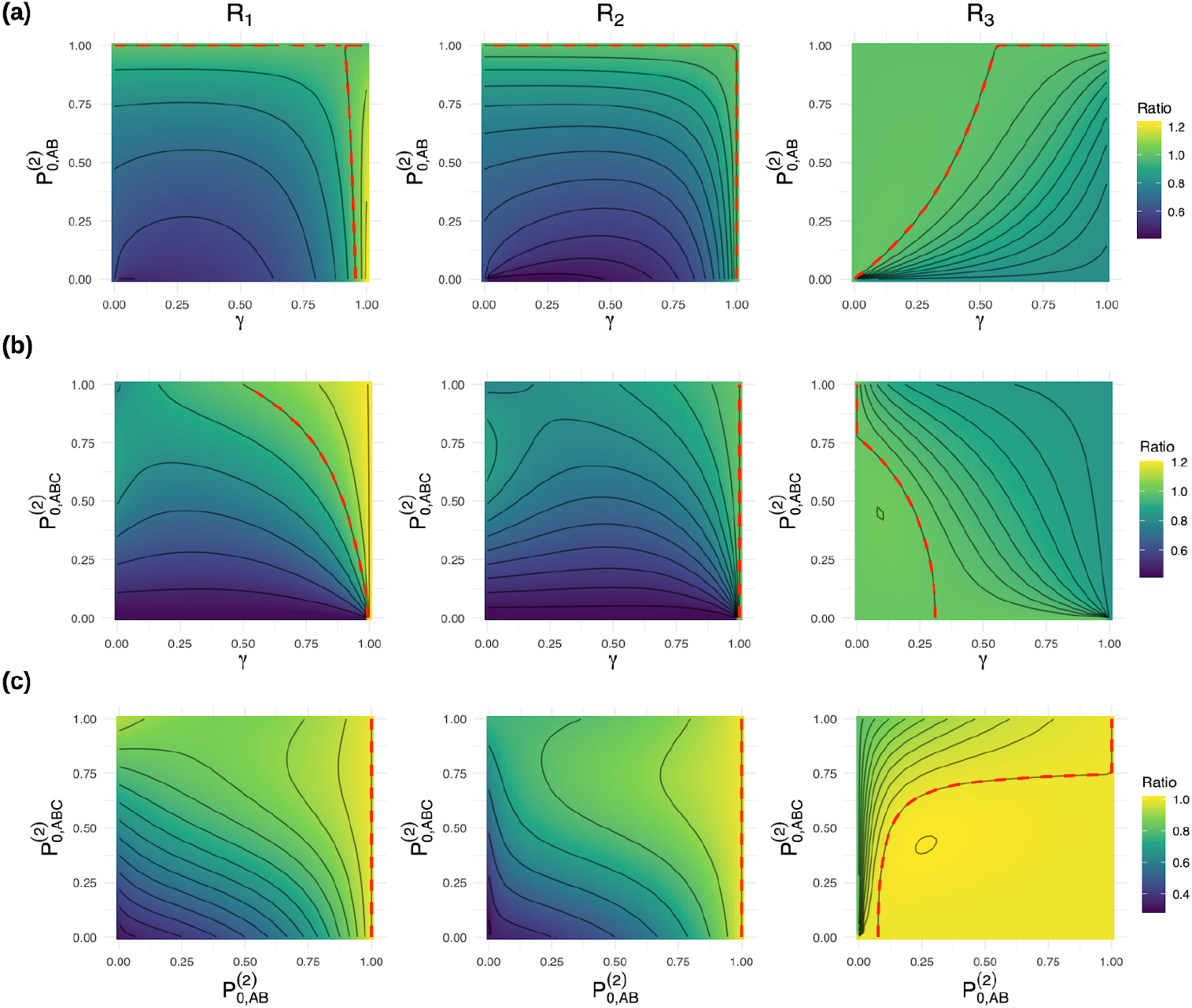
Contour plots of the expectation surface under MSCig model: a)-c) The changes of expectation ratio values in respect to the varying parameters of the MSCig model. The *R*_1_ trough *R*_2_ represent expectation ratios, i.e. 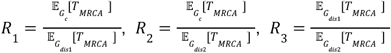 The red dashed line mar*k*s the region where expectation ratios attain unity. *P*^(2)^_0, *AB*_ and *P*^(2)^_0,*ABC*_ denote the probabilities of two lineages not coalescing in the populations *AB* or *ABC* respectively. The γ parameter represents introgression probability. Only two parameters were being varied at a time, while the rest were fixed at γ = 0.1, *P*^(2)^_0, *AB*_ =0.3, *P*^(2)^_0,*ABC*=_ 0. 3,a= 0. 1, *b*= 0. 2,*c*= 0. 3, *d*= 0.4.

**Fig. 5.**
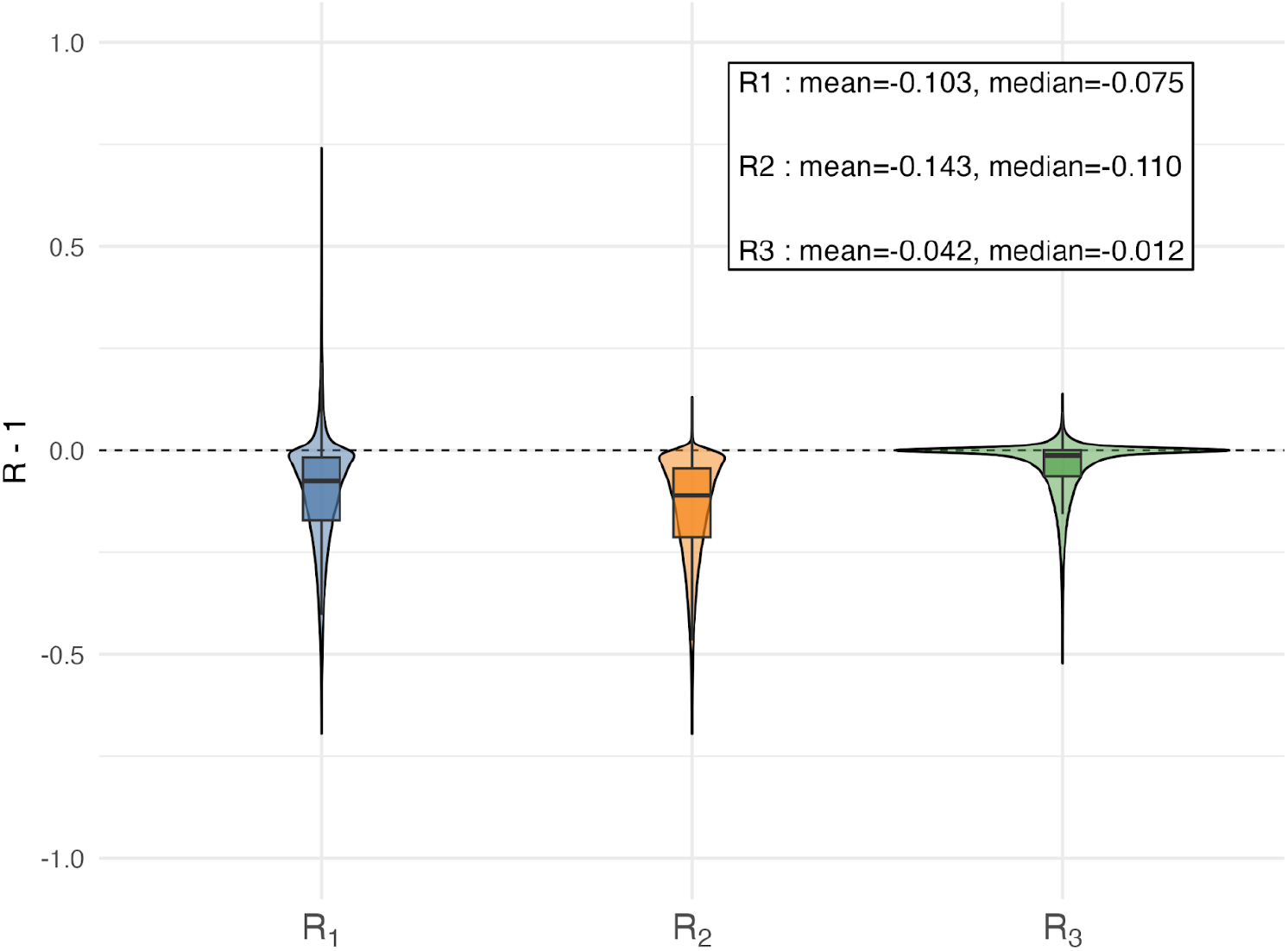
Distributions of simulated expectation ratios under MSCig violin introgression. Violin plots represent simulated distributions of the *R* − 1 values for each of the expectation ratios, i.e. 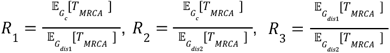.The box plot within each violin shows median as well as lower and upper quantiles. The dashed line mar*k*s the *R*− 1 values of zero.

Finally, to obtain approximations of tree height (*T*_*MRCA*_) for each gene tree triplet we used summary statistic *H*(*T*) which represents the average of tree distances between each tip and the root node, i.e.

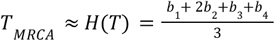

where *b*_1_,*b*_2_, *b*_3_ and *b*_4_ denote branch lengths. The *b*_2_ *b*_3_ and *b*_4_ indicate two sister species and their ancestra branch, respectively, whereas *b*_1_ is the remaining branch that connects to the root node, for example ((*B*: *b*_2_, *C*: *b* _3_):*b*_4_, *A*: *b* _1_). Our statistic *H*(*T*) is related to Sac*k*in index (King and Rosenberg 2021), which is a tree balance statistic representing the sum of all paths from tip to root node on a tree. Thus, *H*(*T*) can be viewed as a normalized Sac*k*in index. By measuring gene tree height as the average distance from all three tips to the root node, (i.e. weighting them equally) we attempt to minimize the bias effect of rate variation between lineages (Fran*k*el and Ané 2023).

### The GhostParser Pipeline

In its core the GhostParser pipeline implements a sequential hypotheses testing procedure (**Fig. 6**) for a rooted species triplet. First, GhostParser compares the frequencies of discordant topologies *P*(*G*_*dis1*_) V *P*(*G*_*dis2*_) using the two proportion Z-test with significance level α = 0.01 (such stringent cutoff is used to reduce the occurrence of false positives), which is a variation of Discordant Count Test (DCT; **Fig. 6**; Suvorov et al. 2022A,B) that is based on χ^2^ test. If this comparison is statistically significantly different then, GhostParser proceeds to comparison of the difference between *H*(*T*) distributions generated from the *G*_*c*_ and *G*_*dis1*_ gene trees (effectively are comparing 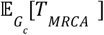 and 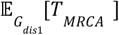), otherwise GhostParser is unable to reject the null hypothesis (i.e., the MSC model), and therefore shows no evidence for introgression. To root each triplet-specific gene tree and calculate *H*(*T*) GhostParser uses ete3 v3.1.3 (Huerta-Cepas et al. 2016) and newic*k*_utils v1.6 (Junier 2024). Then, the resulting *H*(*T*) distributions obtained from a set of *G*_*c*_ and *G*_*dis1*_ gene trees are compared using Tree Height Test (THT) based on the two sample Kolmogrov-Smirnov test with α = 0 05,implemented in R v4.4.2 (R Core Team 2021). If this test does not reject the null hypothesis – i.e. distributions of tree heights from of *G*_*c*_ and *G*_*dis1*_ gene tree sets are similar – GhostParser will suggest sampled introgression has occurred without any evidence of inflow introgression. Further, when THT generates significant result we compare the medians of *H*(*T*) distributions to differentiate between sampled and ghost introgression scenarios. We note that although theoretical expectations were expressed as means, in practice we compare medians of *H*(*T*) distributions, which provide a more robust basis for distinguishing between sampled and ghost introgression. Specifically, if 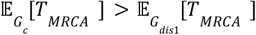 then GhostParser infers sampled inflow introgression (can be with or without outflow), but if 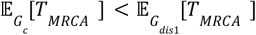 the GhostParser would predict ghost introgression.

**Fig. 6.**
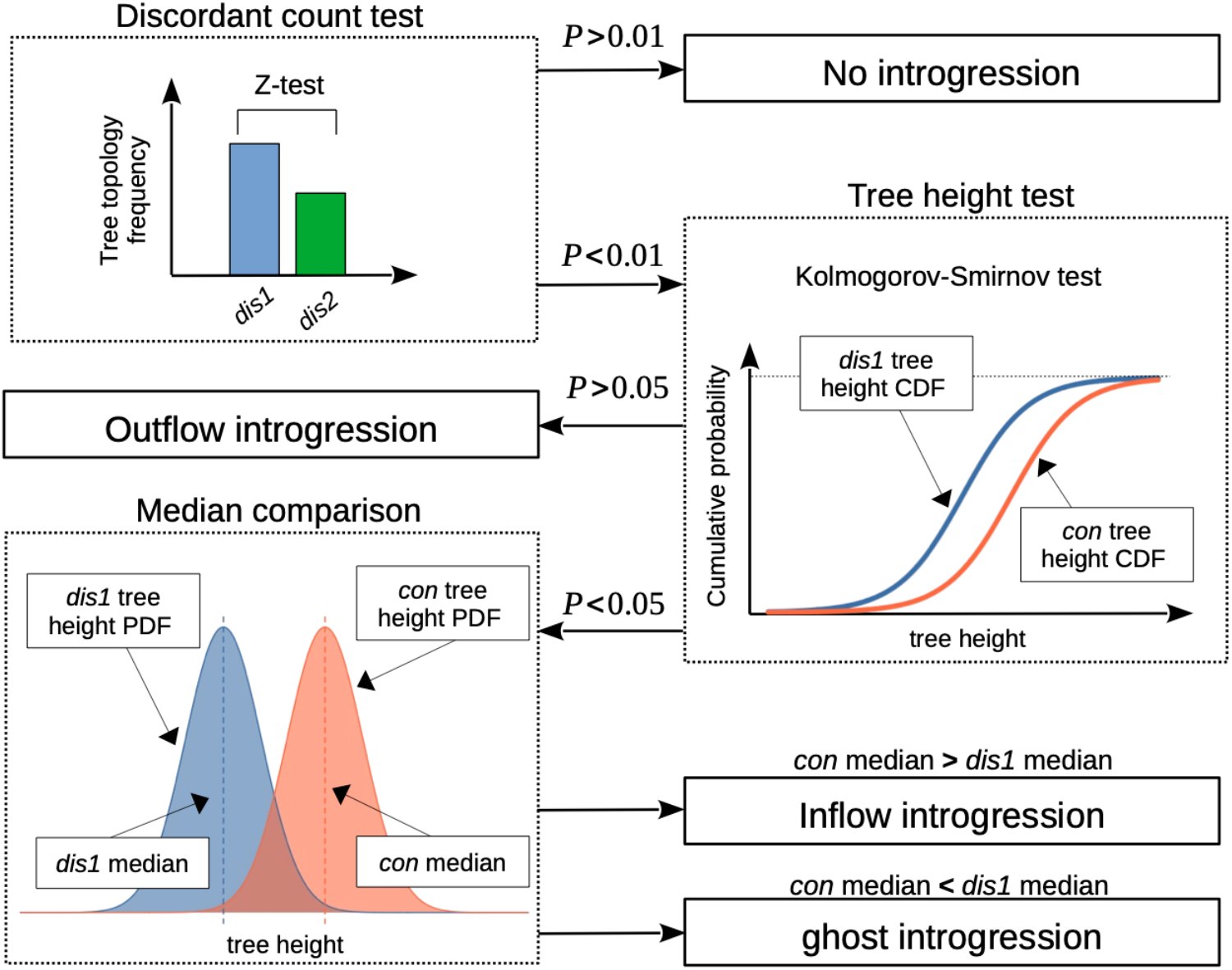
GhostParser pipeline. The GhostParser pipeline follows a sequential hypothesis testing procedure. Key statistical steps are shown in dashed outline boxes, while specific outcomes are shown in solid boxes. PDF probability density function, CDF cumulative distribution function, dis1 discordant topology 1, i.e. ((B,C),A), dis2 discordant topology 2, i.e. ((A,C),B), con = concordant topology, i.e. ((A,B),C).

The GhostParser pipeline represents a python script which uses a combination of python and R languages. In order to launch GhostParser analysis the user provides the following: (i) an input file with gene trees including branch lengths, one per line in newick format, (ii) a text file that contains a outgroup taxa used to root the triplet, and (iii) the rooted species tree topology in newick format. The user may also provide an optional file with the specific triplets of interest for testing. If this optional file is provided, GhostParser will only analyze each user-specified triplet. Otherwise, it will, by default, analyze every possible triplet from the species tree (excluding the triplets that contain an outgroup taxon). GhostParser will classify each of the three taxa in the triplet as the *A B* or *C* taxon. For any species triplet, the *A* and *B* taxa are always defined as sister in concordant gene tree, whereas the *B* taxon is sister to *C* in the most common discordant topology.

GhostFinder is an optional follow-up analysis to GhostParser. It uses the output from GhostParser—which identifies recipients of ghost introgression—to search for the potential “ghost” lineage in the species tree. It identifies, from the GhostParser output file, the triplets where evidence of unsampled introgression has been found. It takes each pairing of *A* and *B* where the *A* taxon is the hypothesized to be a putative recipient of ghost introgression, and the outgroup in *G*_*dis1*_ It then tests all possible *C* taxa from the species tree to determine if any *C* taxa will change the assignments of the sister taxa in the triplet. In instances where the putative ghost recipient is reassigned as the *B* taxon, GhostFinder uses GhostParser to determine which of these *C* taxa show evidence of inflow introgression with the putative ghost, and flag these introgression donors as putative ghost lineages.

### Simulations

Here, we generally followed the simulation approach used by Pang and Zhang (2024) to assess the accuracy of GhostParser, and compared its performance against BPP, which more accurately distinguishes between ghost and sampled introgression than all other methodologies designed to detect both of those introgression scenarios. Since inter-lineage rate variation can generate false positive results when using gene tree summary statistics to detect introgression, we incorporated both inter-lineage and inter-locus rate variation into our simulations to enhance biological realism, following the strategy of Fran*k*el and Ané (2023).

#### Simulating gene trees under differing multispecies coalescent models

To maintain consistency in our simulated dataset and the dataset used to test the accuracy of BPP, we generated our simulated dataset using a modified perl script (`msplus.pl`) from Pang and Zhang’s (2024) work that they used to test the utility of BPP for introgression detection. As originally written, the script uses the *ms* simulator (Hudson 2002) to simulate 480 sets of 1,000 gene trees along a triplet species networ*k* topology. The gene tree sets are generated under ghost, inflow and outflow introgression with specific user-provided introgression probabilities. Specifically, this pipeline generates gene tree sets under eight different combinations of species tree branch lengths, with 20 independent replicates per combination of introgression model and branch lengths. We modified this script to include an additional null model, i.e. MSC, as described below. Here we generated total of 620 sets of 1,000 gene trees each for each of the coalescent models, i.e. MSC, MSCi (outflow), MSCi (inflow) and MSCig.

Each set of gene trees was simulated from a species tree topology of (*X*, ((*A, B*), *C*)) where *X* is an unsampled “ghost” lineage in *ms*. The event times, such as species divergencies and introgressions are specified directly in coalescent units, where 1 unit corresponds to 4*N*_*e*_ generations for a diploid population. We therefore set all event times in these scaled units without specifying *N*_*e*_ explicitly. The distance between MRCA of *ABCX* and MRCA of *ABC* was always set to 5 coalescent units, while the internal branch length of *AB* the distance between the MRCA of *AB* and an introgression event involving either *A* or *B* and the distance between the introgression event and the tree tips are all set at either 0.5 or 5 coalescent units. This results in a total of eight possible branch-length combinations. To simulate aforementioned gene tree sets we used MSC (γ_1_ =γ_2_ =γ =0), MSCi (outflow; γ _1_∈[0 1] γ_2_ =γ =0), MSCi (inflow; γ_2_∈[0 1] γ_1_= γ= 0) and MSCig (γ∈[0 1] γ_1_= γ_2_ =0) models with corresponding introgression probabilities Γ Further, for each possible combination of coalescent model and branch-lengths, 20 independent replicates were generated, for a total of 640 sets of 1000 gene trees each. We produced the simulated gene trees under high (Γ= 0. 3, Γ= 0. 5), moderate (Γ= 0. 1) and low (Γ= 0. 05,Γ= 0.01) values of introgression probabilities, whereΓ =(γ_1_, γ_2_,γ) In summary, this amounted to a total of 3,200 sets of 1,000 gene trees each under every possible combination of introgression probability, timing, and coalescent model including 20 independent replicates.

#### Incorporating rate variation between lineages

To test how GhostParser performs in the presence of rate variation, which can result in false positives of both sampled and ghost introgression, we implemented an approach used by Frankel and Ané (2023) to introduce rate variation into the terminal branches of simulated gene trees. Specifically, for every set of 1,000 gene trees, we first pruned the ghost taxon *X* then we assigned a rate multiplier for each terminal branch in the species tree ((*A*,*B*),*C*) These rate multipliers were drawn independently from a log-normal distribution (μ= 1. 0, σ =0. 7), with a maximum allowed value of four. We then multiplied the branch lengths (expressed in coalescent units) in each of the 1,000 gene trees by the branch rate multiplier.

#### Generating multiple sequence alignments

In generating multiple sequence alignments (MSAs) for each set of 1,000 uniformly rate-modified gene trees, we additionally incorporated intergeneric rate variation, which is observed in empirical multi-locus datasets (Yang 1996; Phillips et al. 2004). We generated each MSA with AliSim (Ly-Trong et al. 2022), implemented in IQ-TREE2 v2.3.6 (Minh et al. 2020). To convert branch lengths from our rate-modified coalescent units to substitutions per site as well as to include intergenic rate variation in each 1,000 gene dataset, we sampled the rate-scaling factor for each locus independently from a Gaussian distribution (μ= 0.036, σ= 0.003). The scaling factor θ 4*N*_*e*_ μ thus, assuming a constant *N*_*e*_ drawing the scaling factor independently from a distribution imbues variation in per-site mutation rate *µ*_*g*._ To maintain consistency with Pang and Zhang’s (2024) simulations, we used the HKY nucleotide substitution model, which assumes unequal transition and transversion rates, and unequal base frequencies (Hasegawa et al. 1985). By default, AliSim assigns transition and transversion rate parameters, as well as base frequencies for the HKY model by drawing corresponding parameters form empirical distributions estimated from a large dataset consisting of different loci (e.g. codon positions, rR 203, Column 3NA, rTNA, introns, intergenic spacers, and UCEs) and genomes (nuclear, mitochondria, virus, and plastid) sampled across the Tree of Life (Naser-Khdour et al. 2025). The alpha parameter for the gamma distribution, which controls rate variation among sites, was drawn from a log-normal distribution, scaled by the the proportion of invariant sites (I) to reflect the relationship between these two parameters 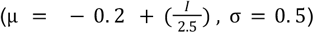 with minimum and maximum constraints of 0.2 and 5.0 to avoid biologically unrealistic values for each gene. To incorporate realistic among-site rate heterogeneity, we modeled the proportion of invariant sites (I) as a random variable drawn from a Beta distribution (α =5, β= 10) which was then scaled by 0.5. These specific parameters produce a right-skewed distribution favoring lower to intermediate proportions of invariant sites, reflecting patterns observed in many empirical datasets where most sites evolve at non-zero rates but some remain highly conserved (Yang 1994; Yang 1996). The additional 0.5 scaling limits invariant sites to at most half the MSA length, maintaining sufficient phylogenetic signal while still allowing considerable conservation. To incorporate insertion and deletion events into our simulations, we drew insertion and deletion rates separately from a Gaussian distribution (μ= 0. 005, σ =0. 00156), with a hard lower bound of zero to prevent the proportions from being assigned negative values. Each simulated MSA contained 1000 sites.

#### Estimating Gene Trees

As errors in inferred MSA can affect gene tree inference (Ogden and Rosenberg 2006; Wong et al. 2008), we realigned every simulated loci with MAFFT v7.525 (Katoh and Standley 2013) with the option *--linsi* to ensure better MSA inferential accuracy. We then estimated maximum likelihood gene tree topologies from the simulated MSAs using IQ-TREE2 v2.3.6 (Minh et al. 2020) under the best fitting substitution model. Finally, we tested GhostParser’s performance on the dataset.

#### Additional testing of potentially problematic demographic scenarios

It has been noted that DCT and BLT based methods could, theoretically, be influenced by ancestral population structure (Suvorov et al. 2022A). Particularly in a scenario where the true species tree is ((*A*,*B*,) *C*) but *B* and *C* share a more recent MRCA, and *A* introgressed with common ancestor of *B* and *C* with a continued gene flow with *B* following the split of *B* and *C* The current understanding is that such a phenomenon may be unlikely (Suvorov et al. 2022A), nonetheless in addition to our *ms* simulated dataset, we generated 500 such scenarios with msprime v1.3.3 (Baumdic*k*er et al. 2022). For each scenario, the MRCA of *ABC* in the species tree was selected from a uniform distribution of between 1 and 50 million generations before present. The *A* lineage and the ancestral branch *BC* each generated gene flow into the other population with the introgression probability drawn from random uniform distribution between 10^-6^ and 0.5 until a random time between the MRCA of *ABC* and 90% of the MRCA of *ABC*, when *B* and *C* diverged. *A* and *B* continued to hybridize, the introgression probability from each population again chosen from a uniform distribution between 10^-6^ and 0.5 until a random time between the MRCA of *BC* and 90% of the MRCA of BC, when gene flow stopped. The *N*_*e*_ of *ABC*, the MRCA of *C* and *B*, and the MRCA of *A, B* and *C* were selected from a uniform distribution *U*(50,000, 5,000,000). The outgroup taxa used to root the triplet was always constrained to diverge with *ABC* 10 million generations prior to the MRCA of *ABC*, and had an *N*_*e*_ of 500,000.

Under each scenario we simulated 1000 gene trees with branch lengths measured in coalescent units. As we only consider the impact of demographic history here, we used the simulated gene trees directly as input for GhostParser testing the hypothesis that *A* was the recipient of ghost introgression or that *B* and *C* introgressed.

### Comparing GhostParser and BPP

We examined the performance of GhostParser on each simulated dataset generated by ms. In contrast to GhostParser, model selection in BPP is computationally intensive—6 CPU-hours per dataset to compare ghost, inflow, and outflow models (Pang & Zhang, 2024). Given this computational cost and our inclusion of an additional no-introgression model, we evaluated BPP only on a subset of simulated datasets. To estimate the performance of BPP in a broad range of scenarios we considered only five of the replicates for each unique combination of timing, introgression probability and coalescent model, instead of the 20 total replicates used to test GhostParser. Additionally, we did not evaluate the performance of BPP in the potentially problematic demographic scenario nor in the scenario of lowest introgression probability (i.e. Γ =0.01). We built the multispecies coalescent with introgression (MSci) scenario to test each of the coalescent models (i.e. MSC, MSCi inflow, MSCi outflow and MSCig) using the “msci-create” function from BPP (see **supplementary table 1** for full commands). We used the same demographic parameters as in Pang and Zhang (2024). Specifically, we used Gaussian quadrature combined with thermodynamic integration to calculate the marginal li*k*elihoods of each scenario. We ran a total of 1.05×10^6^ iterations, using the first 50,000 MCMC iterations as burn-in, and then sampling every other iteration. We then estimated the optimal parameters of the MSci model using the A00 analysis built into BPP (Yang 2015; Pang and Zhang 2024). Each BPP analysis was executed twice to ensure model convergence.

We compared the performance of BPP and GhostParser under each tested level of Γ (with the exception of level of Γ= 0. 01 upon which BPP was not tested). For each tested level of Γ, we calculated the specificity (true negatives/(true negatives + false positives)) and sensitivity (true positives/(true positives + false negatives)) diagnostic metrics of both BPP and GhostParser for our four tested introgression models. We also calculated the overall accuracy for both BPP and GhostParser for each compared level of Γ both when considering inflow and outflow as separate categories, and when considering them as one category of “sampled introgression.”

Since previous study (Pang and Zhang 2024) indicates that BPP exhibits mar*k*edly better performance than other methods designed to differentiate between ghost and sampled introgression (e.g. HyDe (Blischa*k* et al. 2018), SNaQ (Solís-Lemus et al. 2017) and PhyloNet (Li et al. 2022)), we did not consider any other software comparisons.

### Empirical datasets

To further benchmar*k* GhostParser against BPP, we ran GhostParser on two empirical phylogenomic datasets – two plant genera *Jaltomata* and *Thuja* – which were used by BPP in the original study. In each empirical dataset we used previously identified introgression events as guiding hypotheses for GhostParser. Additionally, for each identified reticulation event in each dataset, we calculated the proportion of triplets that support the proposed introgression hypothesis, which can provide a measure of confidence in the inferred introgression event (Suvorov et al., 2022A).

#### Jaltomata

*Jaltomata* is a plant genus of approximately 60 species in the family Solanaceae, which contains subgroups distinguished by mature fruit colors, and has experienced recent radiations (Miller et al. 2011). Using BPP, Pang and Zhang (2024) identified three nodes in the phylogeny of *Jaltomata* influenced by introgression, in a transcriptomic dataset containing 14 species from the genus (Wu et al. 2018). In comparison, node one was incorrectly identified as a recipient of ghost introgression, by HyDe and SNaQ (Solís-Lemus and Ané 2016; Solís-Lemus et al. 2017), whereas BPP correctly identified sampled introgression. The second and third nodes were identified as sampled introgression by BPP, SNaQ and HyDe. The gene trees provided in Pang and Zhang (2024) were used as input for GhostParser to test for introgression in each of the three previously identified nodes.

#### Thuja

Compared to many taxa in the Tree of Life, the patterns of ghost introgression have been extensively categorized in the coniferous plant genus *Thuja* (Li et al., 2022; Pang and Zhang 2024). Li et al., (2022) generated a transcriptomic dataset of 2,969 single-copy genes, for four species of *Thuja*, namely *T. standishii, T. sutchuenensis, T. plicata* and *T. dolobrata*. Of these four species, *T. dolobrata* is the outgroup, while *T. sutchuensis* and *T. standishi* form sister relationships (Li et al. 2022). Li et al., (2022) identified *T. sutchuenensis* as the recipient of ghost introgression, while HyDe identified introgression between *T. standishi* and *T. plicata* (Pang and Zhang 2024). Pang and Zhang (2024), used BPP to tease apart the competing hypotheses that *T. standishi* and *T. plicata* experienced gene flow, whereas *T. sutchuenensis* was found to be the recipient of ghost introgression. As with the *Jaltomata* dataset, the *Thuja* gene trees provided by Pang and Zhang (2024) were used as the input for GhostParser.

### Additional empirical testing of GhostParser

To further demonstrate the utility of GhostParser in empirical systems we tested GhostParser in two clades which are now common examples of widespread introgression. First, identification of introgression across the butterfly genus *Heliconius* was one of the first discoveries to support the hypothesis that introgression played a key role in the radiation of insects (Edelman et al. 2019). Second, the fly genus *Drosophila* exhibits abundant instances of introgression despite evidence for reproductive isolation (Suvorov, et al. 2022A). However, both studies relied on introgression estimation methods that are incapable of identifying ghost introgression and are prone to producing false positive results in the presence of inter-lineage rate variation (Fran*k*el and Ané 2023). Here, we use GhostParser to revisit the hypotheses of introgression identified in these systems.

#### Heliconius

Here, we revisit the hypothesis of sampled introgression in the melpomene clade of *Heliconius* butterflies, using trees generated from a whole genome alignment of *Heliconius* (Edelman et al. 2019). We reconsider two introgression events with GhostParser using the original set of gene trees as input, and using triplets from two hypothesized introgression events as guides for hypothesis testing in this classical system, with the tree topology previously estimated by Edelman et al., (2019) as GhostParser input.

### Drosophila

The identification of introgression in the fly genus *Drosophila* is among the most promising systems for exploring the functional impacts of introgression, as the genomes of the model species *D. melanogaster* and *D. simulans* are amongst the most well studied in the entire Tree of Life (Suvorov, et al. 2022A). Here, we considered the *Drosophila* gene tree dataset, generated by Suvorov et al. (2022A). The original analysis of this dataset identified numerous putative instances of introgression. In particular, clades three, seven and nine contained two, five and three instances of highly supported inter-specific gene flow respectively. We used this dataset to revisit each of these putative introgression events with GhostParser, and tested an additional six species triplets where no introgression was detected to test for putative false positives. In detected instances of ghost introgression, we employed GhostFinder to search the *Drosophila* tree for possible “ghost” lineages across the entire sampled *Drosophila* dataset, using the species tree estimated by Suvorov et al., (2022A) as input.

## Results

### Comparison of GhostParser and BPP in simulated datasets

The model accuracy of GhostParser was greater than that of BPP both when inflow and outflow introgression were considered to be two separate categories, and when they were combined into one category of sampled introgression, regardless of the introgression probability tested (**Table 1; Fig. 7**). Both BPP and GhostParser maintained similar accuracy under Γ =0.50 (GhostParser = 0.91, BPP = 0.71 with inflow and outflow introgression combined) and Γ =0.30 (GhostParser = 0.92, BPP = 0.69 with inflow and outflow introgression binned together; **Table 1**).

**Table 1:** Comparison of GhostParser and BPP performance Sensitivity and specificity for each introgression model are shown for both BPP and GhostParser under tested introgression probabilities (Γ). BPP was not tested at Γ=0.01. MSC = multispecies coalescent, MSCi = multispecies c alescent with introgression, MSCig = multispecies coalescent with ghost introgression.

| **Table 1: Comparison of GhostParser and BPP performance** |  |  |  |  |  |  |  |  |  |  |
| --- | --- | --- | --- | --- | --- | --- | --- | --- | --- | --- |
|  | MSCig<br>sensitivity | MSCig<br>specificity | MSCi<br>(inflow)<br>sensitivity | MSCi<br>(inflow)<br>specificity | MSCi<br>(outflow)<br>sensitivity | MSCi<br>(outflow)<br>specificity | MSC<br>sensitivity | MSC<br>specificity | Overall<br>model<br>accuracy<br>(inflow<br>and<br>outflow<br>separate) | Overall<br>model<br>accuracy<br>(inflow<br>and<br>outflow<br>combined) |
| **$\Gamma = 0.50$** |  |  |  |  |  |  |  |  |  |  |
| GhostParser | 0.93 | 0.91 | 0.88 | 0.86 | 0.37 | 0.95 | 0.99 | 1.00 | 0.79 | 0.91 |
| BPP | 0.85 | 0.91 | 0.97 | 0.57 | 0.52 | 0.98 | 0.07 | 0.76 | 0.61 | 0.71 |
| **$\Gamma = 0.30$** |  |  |  |  |  |  |  |  |  |  |
| GhostParser | 0.91 | 0.95 | 0.88 | 0.91 | 0.44 | 0.95 | 0.97 | 0.99 | 0.80 | 0.92 |
| BPP | 0.82 | 0.88 | 0.90 | 0.58 | 0.50 | 1.00 | 0.10 | 0.77 | 0.58 | 0.69 |
| **$\Gamma = 0.10$** |  |  |  |  |  |  |  |  |  |  |
| GhostParser | 0.67 | 0.99 | 0.61 | 0.81 | 0.37 | 0.92 | 0.98 | 0.84 | 0.66 | 0.80 |
| BPP | 0.50 | 0.92 | 0.87 | 0.47 | 0.52 | 1.00 | 0.12 | 1.00 | 0.50 | 0.61 |
| **$\Gamma = 0.05$** |  |  |  |  |  |  |  |  |  |  |
| GhostParser | 0.35 | 1.00 | 0.41 | 0.82 | 0.22 | 0.96 | 0.98 | 0.55 | 0.50 | 0.60 |
| BPP | 0.38 | 0.90 | 0.80 | 0.39 | 0.45 | 0.98 | 0.05 | 1.00 | 0.41 | 0.53 |
| **$\Gamma = 0.01$** |  |  |  |  |  |  |  |  |  |  |
| GhostParser | 0.03 | 1.00 | 0.03 | 0.98 | 0.02 | 0.99 | 0.99 | 0.06 | 0.27 | 0.24 |
| **Table 1:** Sensitivity and specificity for each introgression model are shown for both BPP and GhostParser under tested introgression probabilities ($\Gamma$). BPP was not tested at $\Gamma = 0.01$. MSC = multispecies coalescent, MSCi = multispecies coalescent with introgression, MSCig = multispecies coalescent with ghost introgression. | | | | | | | | | | |

**Fig. 7.**
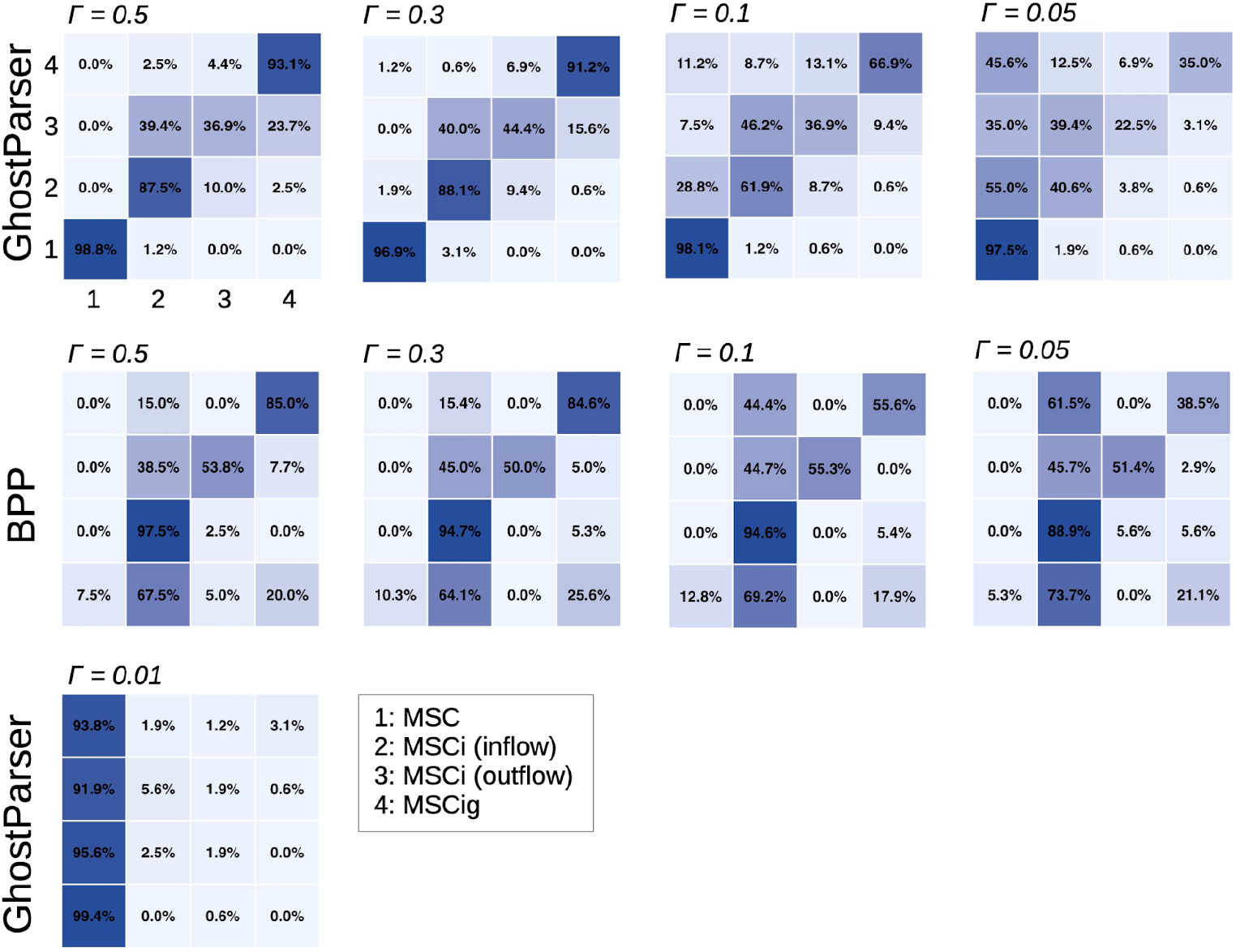
Performance comparison of GhostParser and BPP on simulated datasets. Confusion matrices show percentage accuracy for each class, where Γ indicates introgression probability. Correct classifications appear along the diagonal, while all off-diagonal entries represent misclassifications. MSC multispecies coalescent, MSCi multispecies coalescent with introgression, MSCig = multispecies coalescent with ghost introgression.

Accuracy for both diminished at Γ= 0.1 (GhostParser = 0.80, BPP = 0.61 with inflow and outflow introgression binned together) and Γ= 0.05 (GhostParser = 0.60, BPP = 0.53 with inflow and outflow introgression combined; Table 1). GhostParser performed at random (accuracy = 0.24) under Γ= 0.01 (BPP was not tested under this introgression probability due to computational constraints; **Table 1; Fig. 7**). GhostParser was more sensitive to ghost introgression under Γ= 0.50 and Γ= 0.30 while BPP was more sensitive under the lower strength introgression scenarios (**Table 1**). The specificity of ghost introgression was high (0.88) n both BPP and GhostParser (**Table 1**) under all introgression strengths tested, with GhostParser marginally (difference < 0.10) outperforming BPP in the specificity of ghost introgression, excepting Γ= 0. 50 where both were tied at 0.91 (**Table 1**). BPP was more sensitive to both inflow and outflow introgression than GhostParser (**Table 1**). GhostParser was much more specific (>0.80) in its detection of inflow introgression than BPP (<0.60). BPP showed higher sensitivity to outflow introgression than GhostParser (**Table 1**). Both were highly specific in their detection of outflow introgression (0.95) with BPP only outperforming GhostParser by a few points. BPP showed low (<0.15) sensitivity to the detection of no introgression, while GhostParser exhibited very high (>0.97) sensitivity to this model (**Table 1**). BPP was highly specific (>0.99) in its detection of no introgression at Γ =0.50 and Γ =0.30 The specificity of GhostParser under MSC dropped to 0.84 at Γ =0.1 to 0.55 at Γ= 0.05 and to 0.06 at Γ =0.01 (Table 1). BPP showed the inverse relationship with the specificity of the detection of the null model, climbing from 0.76 at Γ =0.50 to 1.00 at Γ= 0.10 (**Table 1**).

### Additional testing of GhostParser in simulated datasets

In the 500 tested replicates of the continuous gene flow scenario (simulated with msprime to test f population structures could produce false positives in GhostParser. GhostParser found no evidence for introgression in 96.6% of the simulated triplets and classified 2.0% and 1.4% of the simulated triplets as sampled and ghost introgression respectively.

### Empirical comparisons to BPP

#### Jaltomata

In scenario in the *Jaltomata* tree, which SNaQ and HyDe identified as ghost introgression, but BPP classified as sampled introgression (Pang and Zhang 2024), all thirty triplets tested with GhostParser supported the hypothesis of sampled introgression (**Fig. 8**). GhostParser classified scenario two (1/1 triplets) as ghost introgression, contrary to BPP, HyDe and SNaQ, and classified scenario three (56/70 triplets) as sampled introgression, as did BPP, HyDe and SNaQ. GhostParser classified each sampled introgression event as “inflow”, contrary to BPP which classified the events as “outflow.”

**Fig. 8.**
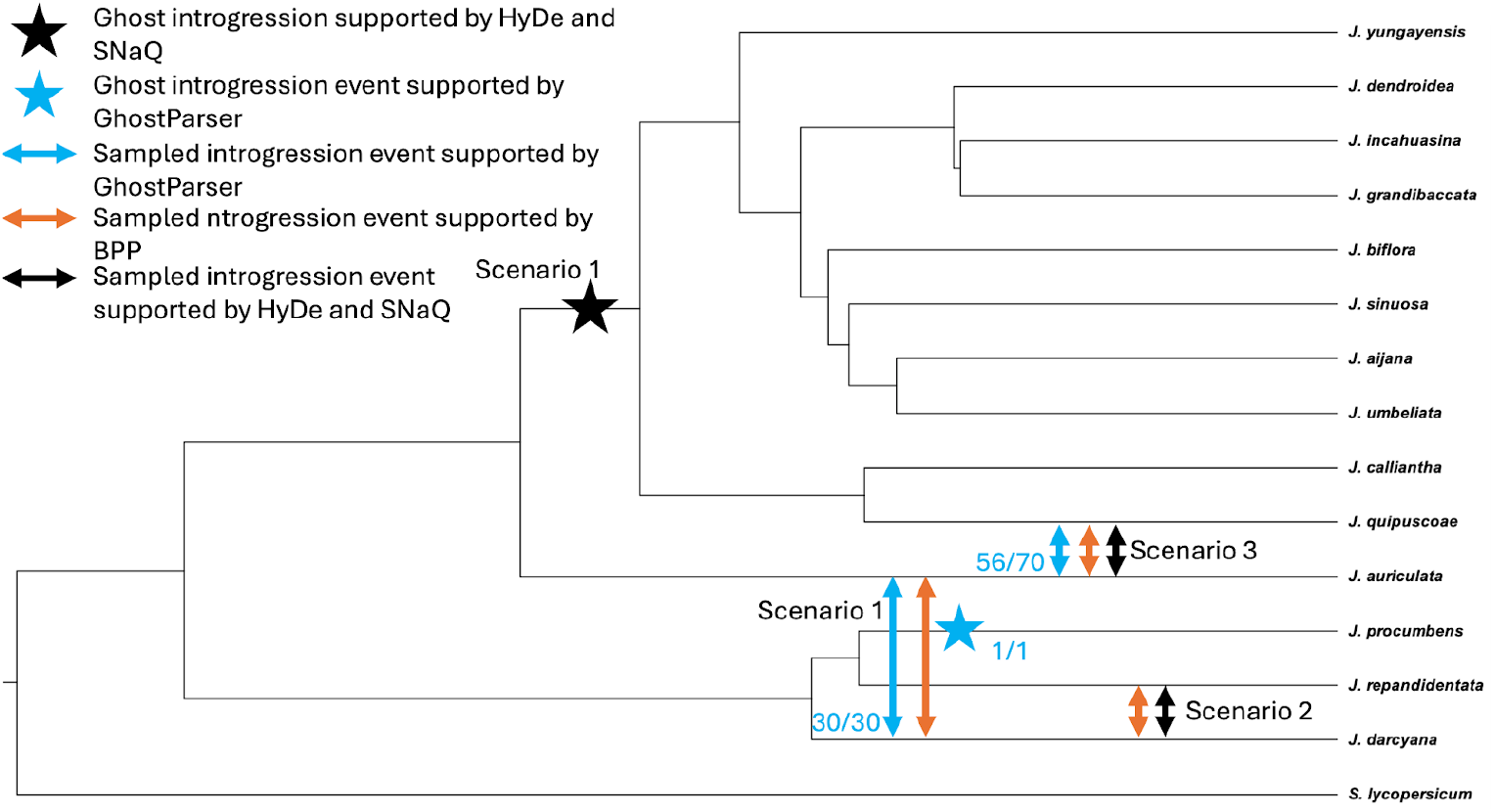
Introgression within the genus Jaltomata: Scenario is a reticulation event classified as ghost introgression by summary statistic based method HyDe and the networ*k* method SNaQ, but as sampled introgression by the full li*k*elihood method BPP (Pang and Zhang 2024) and GhostParser. Scenarios two and three wer classified as sampled introgression by HyDe, SNaQ, BPP (Pang and Zhang 2024), while only scenario three wa classified as sampled introgression by GhostParser. The ratios represent the proportion of triplets supporting the introgression hypothesis (either ghost or sampled) in GhostParser.

#### Thuja

In the species triplet ((*T. sutchuensis, T. standishi*), *T. plicata*) GhostParser supported the hypothesis of *T. sutcheuensis* being the recipient of ghost introgression, a result consistent with BPP and PhyloNet findings, whereas HyDe inferred only sampled introgression between *T. plicata and T. standishi.*

#### Heliconius

GhostParser identified both instances of introgression identified by QuIBL in the melpomene clade of *Heliconius* as ghost introgression, with the *H. cydno, H. timareta, H. melpomene* clade identified as the recipient of ghost introgression (**Fig. 9**). However, in all tested triplets, the gene trees supporting the sampled introgression hypothesis (from QuIBL) had a higher count than the species tree topology (Fig. 9), suggesting the species tree hypothesis we used as input for GhostParser (Edelman et al. 2019) may be incorrect.

**Fig. 9.**
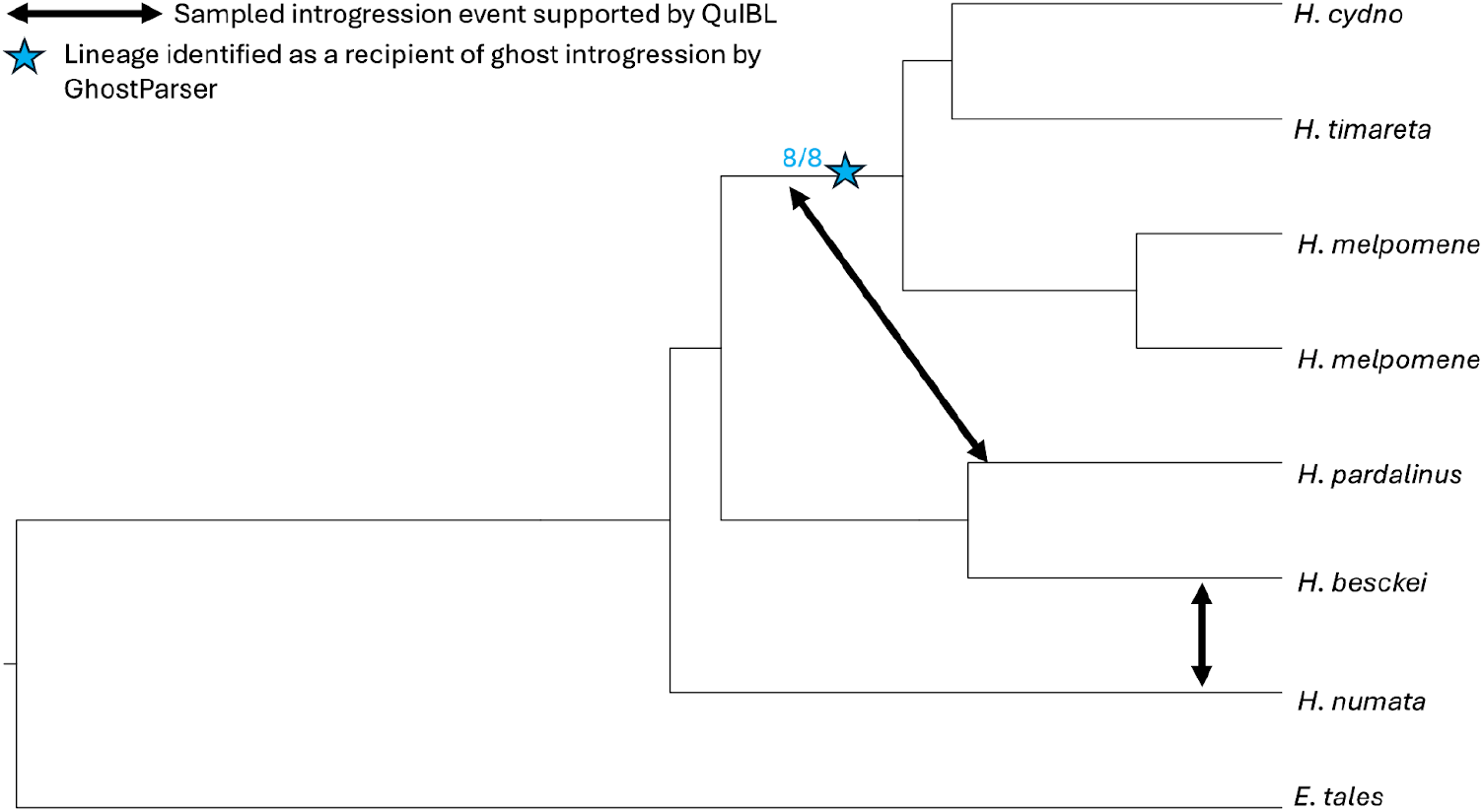
Introgression within the melpomene clade of Heliconious generated from trees of sliding windows of the Heliconius genome alignment (Edelman et al. 2019) Introgression events detected with QuIBL are shown in blac*k* arrows, and ghost introgression detected with GhostParser are shown with blue stars alongside the number of supporting triplets.

#### Drosophila

One of the sampled introgression events in Clade 2 of *Drosophila* was classified as ghost introgression (7/9 triplets) by GhostParser, while the other gene flow events showed support for sampled introgression (25/28; **Fig. 10**). Two events in Clade 7 were classified as sampled introgression by GhostParser (**Fig. 10**), with the other two being classified as ghost introgression. One of the introgression events in Clade 9 showed mixed support for sampled (60/110 triplets) and ghost (50/110) introgression, while the other two were classified as sampled introgression (**Fig. 10**). Ghostfinder did not identify any other taxa in the species tree which introgressed with the ghost recipients in either Clades 2 or 7. This suggests that the “ghost” lineages were li*k*ely unsampled sister groups to Clades 2 and 7 respectively.

**Fig. 10.**
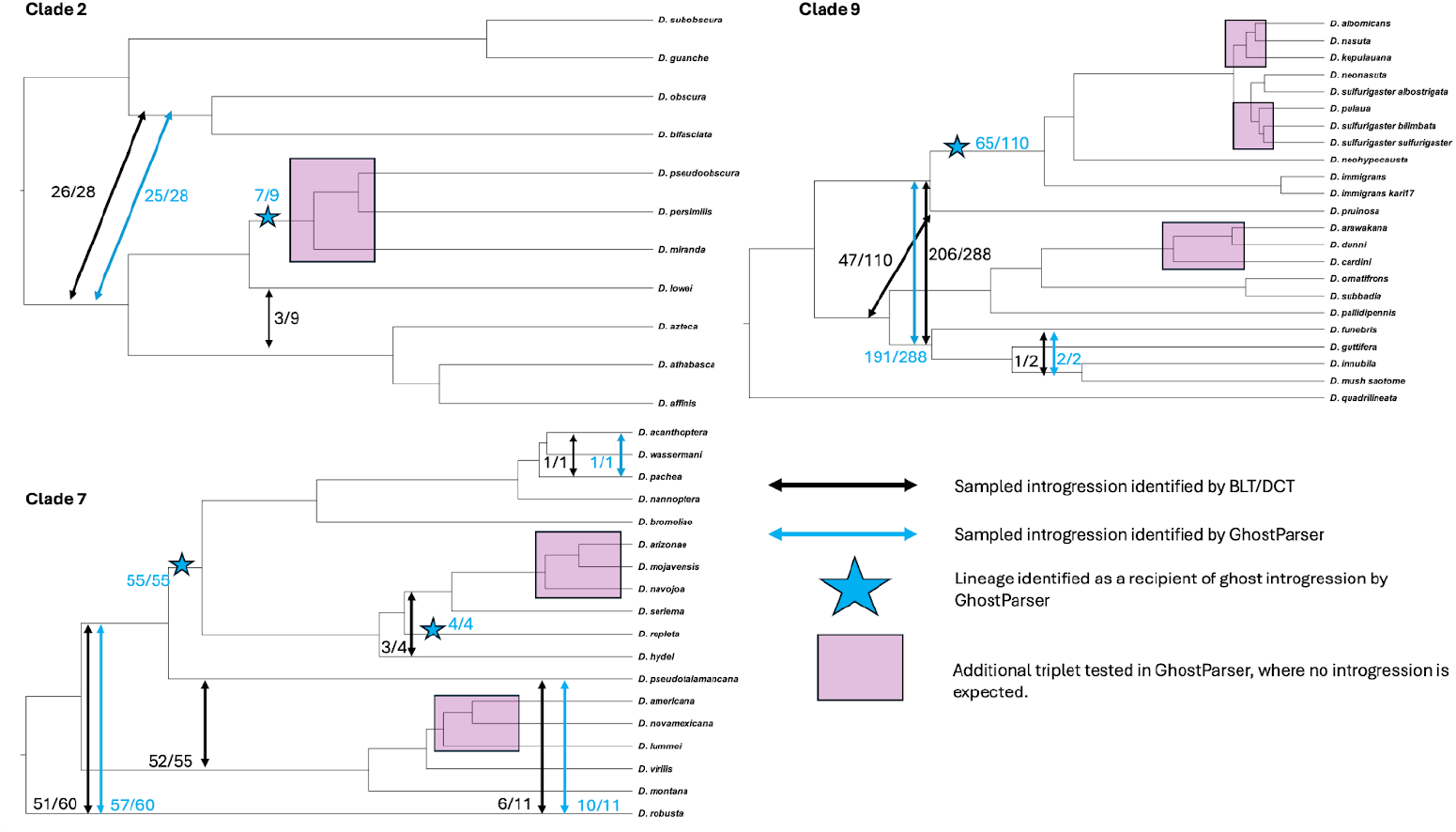
Introgression across clades 2, 7, and 9 of Drosophila: Previously detected introgression events with BLT/DCT shown in black arrows, with the number of supporting triplets in blac*k*. Sampled introgression events detected by GhostParser are shown in blue arrows with the number of supporting triplets in blue, and lineages identified as recipients of ghost introgression are marked with blue stars, with the number of triplets supporting ghost introgression also shown in blue text. Additional clades tested with GhostParser are highlighted in pink. GhostParser did not identify reticulation events in any of these clades. The proportion of triplets supporting the introgression hypotheses from BLT and DCT are shown in black, while the proportion of triplets supporting the GhostParser hypothesis are shown in blue.

## Discussion

### Towards scalable selection of an appropriate introgression model in the presence of rate variation

Despite the major influence that both sampled and ghost introgression can have on evolutionary inference (Whitney et al. 2006; Pardo-Diaz et al. 2012; Huerta-Sánchez et al. 2014; Bastide et al. 2018; Jones et al. 2018; Taylor and Larson 2019; Ottenburghs 2020b; Gibson et al. 2021; Hibbins and Hahn 2021), no bioinformatic tools exist which can accurately, robustly and in a scalable manner distinguish between these two introgression scenarios from multi-locus phylogenomic datasets (Pang and Zhang, 2024), particularly in the presence of between lineage rate variation (Frankel and Ané 2023; Koppetsch et al. 2024). By considering a novel summary statistic *H*(*T*) used by our THT, we successfully identify the presence and absence of ghost and sampled introgression in simulated datasets.

Indeed both simulated and empirical results provide compelling evidence that GhostParser is a reliable approach to estimate both ghost and sampled introgression, even when only one triplet is available for testing, as was the case in the simulated datasets. Not only is the sensitivity of GhostParser to ghost introgression high (>75%) when Γ ≥ 0.1 in our simulations, but GhostParser also rarely misclassifies cases generated under either MSC or MSCi models as ghost introgression (**Table 1**). Nor did GhostParser identify introgression in the tested empirical triplets for which other methodologies did not identify evidence of introgression (**Fig. 10**). GhostParser concurred with BPP in the empirical *Jaltomata* and *Thuja* datasets, even at nodes where other network and summary-statistic based tests differed, with the exception of classifying one reticulation event in *Jaltomata* as ghost introgression (**Fig. 8**). However, the overall accuracy of GhostParser drops when inflow and outflow are considered to be two separate categories, as does the accuracy of the state-of-the-art method BPP (**Table 1**).

BPP exhibited inferior performance in comparison with GhostParser in terms of accuracy criteria metrics (**Table 1**). In particular, BPP produced less accurate results (overall accuracy) than GhostParser, failing to distinguish between simple MSC models with no introgression and MSCi models regardless of Γ, and in distinguishing between inflow and outflow introgression (**Table 1**). We do note that Pang and Zhang’s (2024) work did not include a no-introgression scenario in their initial tests of the use of BPP to distinguish between ghost and sampled introgression hypotheses. Furthermore, unlike BPP, GhostParser was highly scalable. It considered each tested triplet in <20 seconds, compared to the hours, or even days, of real time required by the full-likelihood method employed by BPP. This makes GhostParser more accessible to researchers with more limited computational resources.

According to our current understanding of introgression and rate variation, GhostParser succeeds where other heuristic and pseudo-likelihood tests have failed because of its usage of the *H*(*T*) statistic in its THT approach. Other branch length tests compare summaries of distances between sister taxa.. For example, The BLT method will compare the distribution of genetic distances between the sister pair *BC* in *G*_*dis1*_ and *AC* in *G*_*dis2*_ if these distances significantly differ, *B* and *C* are assumed to have introgressed (Suvorov et al., 2022A). If, for example, *B* evolved more rapidly than *C* these genetic distances will be skewed mimic*k*ing an introgression signal (Frankel and Ané 2023). Likewise, if *A* introgressed with the ghost lineage *X* the proportions of *G*_*dis1*_ and *G*_*dis2*_ would be significantly different, causing the DCT to infer introgression between *B* and *C* (Suvorov et al., 2022A), while the BLT concludes no introgression has occurred. By using the *H*(*T*) statistic, GhostParser generates a low false positive rate for introgression detection in the presence of between lineage rate variation allowing reliable inference of sampled and ghost introgression.

### Suggested Implementation of GhostParser

GhostParser can be implemented as a stand alone tool to detect introgression, or as a follow up analysis to assess results of prior introgression estimates produced by various methods. GhostParser can be used to test specific hypotheses, and determine if the identified gene flow events may be the result of ghost introgression, or if introgression has occurred at all. While the sensitivity and accuracy of GhostParser was diminished for simulated scenarios with low introgression probability (i.e. Γ < 0.1), as compared to larger tested values of introgression strength, this did not appear to hamper its ability to detect ghost and sampled introgression in sampled empirical systems. Nevertheless, BPP is marginally more sensitive to ghost introgression at Γ 0 05 thus the user may want to consider further analysis with BPP if they suspect a false negative result. Under all scenarios of introgression strength (Γ= 0.01, 0.05, 0.10,0.30, 0.50), GhostParser remains highly specific in its classification of ghost introgression (**Table 1**). We do caution against relying on GhostParser or BPP to identify the presence of inflow. Accordingly, we did not consider inflow in our empirical dataset analyzed with GhostParser, except in systems where BPP was explicitly used to model the directionality of gene flow. The GhostParser approach distinguishing between ghost and sampled introgression opens the field of reliable and fast introgression detection to those limited by time and computational resources. This will be an important software in introgression detection and systematics.

## Data Availability

The ghostparser software, tutorial, and all data for the empirical systems tested are publicly available on github (https://github.com/asuvorovlab/ghostparser.git). The simulated dataset used to test the accuracy of GhostParser, including the gene trees, gene tree alignments, estimated gene trees, and GhostParser output are available on figshare (https://doi.org/10.6084/m9.figshare.29921969).

## Acknowledgements

We would like to thank Daniel Schrider for his helpful feedback on this project. All computational work was conducted on the Advanced Research Computing clusters at Virginia Tech.

## Appendix: Derivation of gene tree *T*_*MRCA*_ expectations under different multispecies coalescent models.

Here, we provide a detailed derivation of three taxon gene tree expectations (𝔼 [*T*_*MRCA*_]) for the multispecies coalescent (MSC), multispecies coalescent with sampled bidirectional introgression (MSCi) and multispecies coalescent with unsampled (“ghost”) introgression (MSCig) models (**Fig 1a-c**). Under all these models 𝔼 [*T* _*MRCA*_] can be viewed as a sum of tree height expectations multiplied by their corresponding probabilities ω For instance, in the MSC model, a concordant gene tree can occur in two different ways, where *A* and*B* lineages either coalesce in the ancestral population *AB* or *ABC*. Each of those two scenarios will have its own probabilities of occurrence ω_1_ and ω_2_ Also, note that ω_1_ + ω_2_ =*P*(*G*_*c*_). Two discordant topologies can only occur in ancestral population *ABC* with probabilities ω_3_ and ω_4_. Additionally, for concordant topology where *A* and *B* coalesce in population *AB* the *T*_*MRCA*_ expectation is *a* 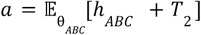,whereas for the other concordant topology and both discordant topologies the *T*_*MRCA*_ expectation is 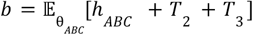. Thus:

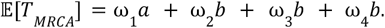

However this model is overparametrized since ω _1_=1 − *P*^(2)^_0,*AB*_ and 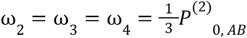, where 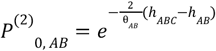. Hence, we can simplify the expectation mixture:

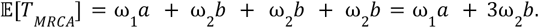

Then, gene tree *T*_*MRCA*_ expectations for concordant and discordant topologies can be obtained by reweighting (normalizing) each individual mixture component, i.e.

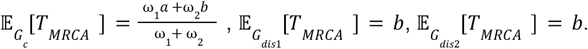

Similarly, we derive total for 𝔼[*T*_MRCA_] MSCi, where additional outflow and inflow Г= (γ _1_,γ_2_)introgression probability parameters are introduced:

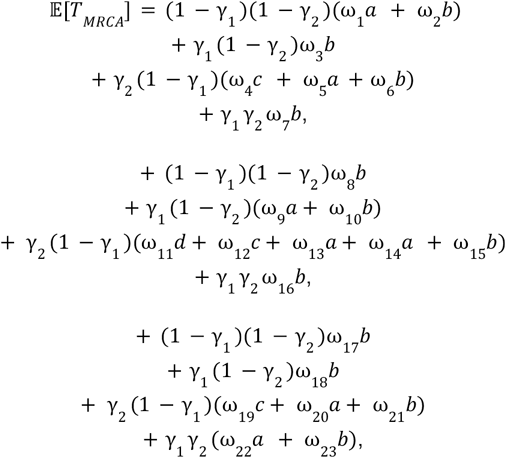

where 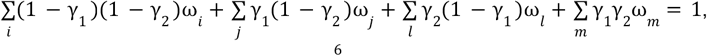 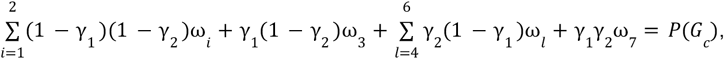 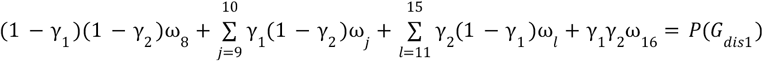 and 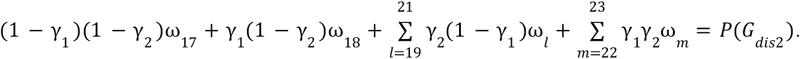.

Here we define three probabilities (i) *P*^(3)^_2, *AB*_ of observing exactly two coalescences for three lineages in a population *AB*, (ii) *P*^(3)^_1, *AB*_ of exactly one coalescence for three lineages in a population *AB* and (iii) *P*^(3)^_0, *AB*_ of no coalescences for three lineages in population *AB* (Rosenberg 2002). Also, note that *P*^(3)^_2 *AB*_ + *P*^(3)^_1 *AB*_ + *P*^(3)^_0 *AB*_ =1.

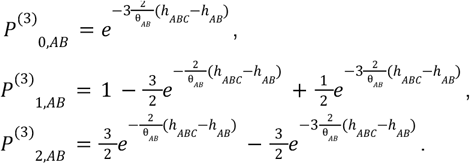

These probabilities that involve coalescence of three branches can be conveniently expressed in terms of 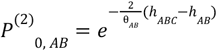, i.e.

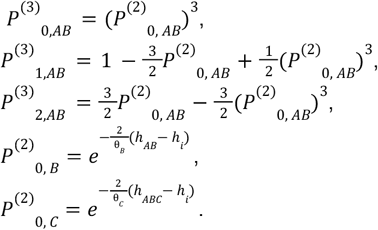

Now, we can establish the equality constraints between different model weights:

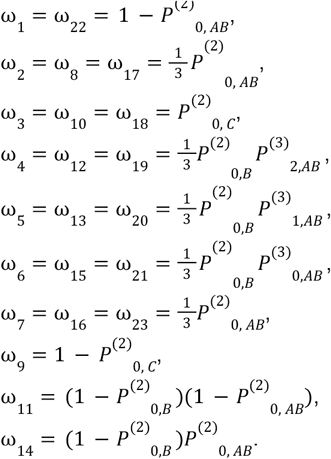

The only weights that are not constrained include (ω_9_ ω_11 14_) and are unique to discordant topology *G*_*dis1*_ mixture component, thus:

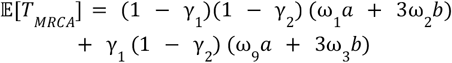

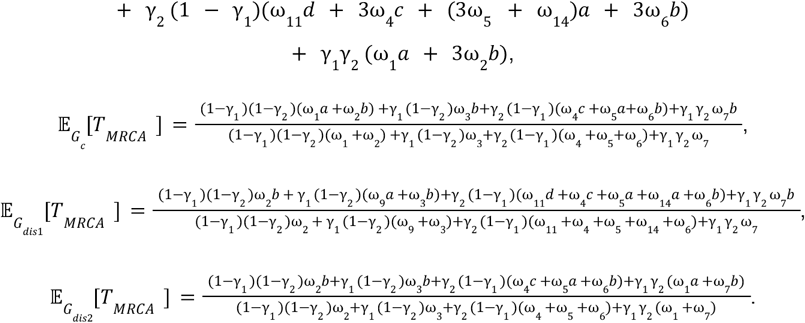

Under MSCig model 𝔼 [*T*_*MRCA*_] is expressed as follows

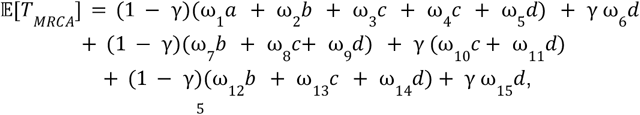

Where 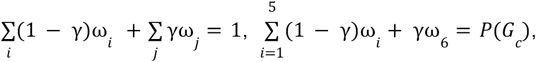 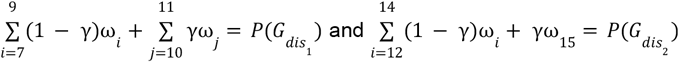, and coalescence probabilities:

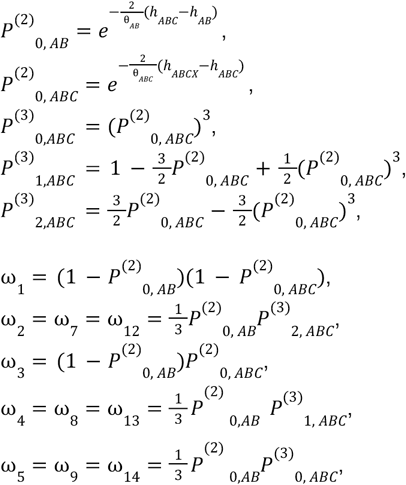

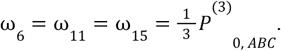

Finally, the applying the weight constraints and mixture weight normalization yields:

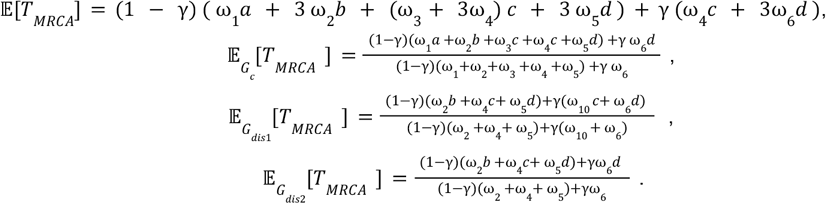

