## Supplementary Material for "GhostParser: A highly scalable phylogenomic approach for the identification of ghost introgression"

### Appendix: Derivation of gene tree $T_{MRCA}$ expectations under different multispecies coalescent models.

Here, we provide a detailed derivation of three taxon gene tree expectations ($\mathbb{E}[T_{MRCA}]$) for the multispecies coalescent (MSC), multispecies coalescent with sampled bidirectional introgression (MSCi) and multispecies coalescent with unsampled ("ghost") introgression (MSCig) models (**Fig 1 a-c**). Under all these models $\mathbb{E}[T_{MRCA}]$ can be viewed as a sum of tree height expectations multiplied by their corresponding probabilities ω. For instance, in the MSC model, a concordant gene tree can occur in two different ways, where $A$ and $B$ lineages either coalesce in the ancestral population $AB$ or $ABC$. Each of those two scenarios will have its own probabilities of occurrence $\omega_1$ and $\omega_2$. Also, note that $\omega_1 + \omega_2 = P(G_c)$. Two discordant topologies can only occur in ancestral population $ABC$ with probabilities $\omega_3$ and $\omega_4$. Additionally, for concordant topology where $A$ and $B$ coalesce in population $AB$ the $T_{MRCA}$ expectation is $a = \mathbb{E}_{\theta_{ABC}}[h_{ABC} + T_2]$, whereas for the other concordant topology and both discordant topologies the $T_{MRCA}$ expectation is $b = \mathbb{E}_{\theta_{ABC}}[h_{ABC} + T_2 + T_3]$. Thus:

$$\mathbb{E}[T_{MRCA}] = \omega_1 a + \omega_2 b + \omega_3 b + \omega_4 b.$$

However this model is overparametrized since $\omega_1 = 1 - P^{(2)}_{0,AB}$ and $\omega_2 = \omega_3 = \omega_4 = \frac{1}{3}P^{(2)}_{0,AB}$, where $P^{(2)}_{0,AB} = e^{-\frac{2}{\theta_{AB}}(h_{ABC} - h_{AB})}$. Hence, we can simplify the expectation mixture:

$$\mathbb{E}[T_{MRCA}] = \omega_1 a + \omega_2 b + \omega_2 b + \omega_2 b = \omega_1 a + 3\omega_2 b.$$

Then, gene tree $T_{MRCA}$ expectations for concordant and discordant topologies can be obtained by reweighting (normalizing) each individual mixture component, i.e.

$$\mathbb{E}_{G_c}[T_{MRCA}] = \frac{\omega_1 a + \omega_2 b}{\omega_1 + \omega_2}, \mathbb{E}_{G_{dis1}}[T_{MRCA}] = b, \mathbb{E}_{G_{dis2}}[T_{MRCA}] = b.$$

Similarly, we derive total $\mathbb{E}[T_{\mathrm{MRCA}}]$ for MSCi, where additional outflow and inflow $\Gamma = (\gamma_1, \gamma_2)$ introgression probability parameters are introduced:

$$\begin{aligned}\mathbb{E}[T_{MRCA}] = &(1 - \gamma_1)(1 - \gamma_2)(\omega_1 a + \omega_2 b)\\ &+ \gamma_1(1 - \gamma_2)\omega_3 b\\ &+ \gamma_2(1 - \gamma_1)(\omega_4 c + \omega_5 a + \omega_6 b)\\ &+ \gamma_1\gamma_2\omega_7 b,\\ &+ (1 - \gamma_1)(1 - \gamma_2)\omega_8 b\end{aligned}$$

$$+ \gamma_1 (1 - \gamma_2)(\omega_9 a + \omega_{10} b)$$
$$+ \gamma_2 (1 - \gamma_1)(\omega_{11} d + \omega_{12} c + \omega_{13} a + \omega_{14} a + \omega_{15} b)$$
$$+ \gamma_1 \gamma_2 \omega_{16} b,$$

$$+ (1 - \gamma_1)(1 - \gamma_2)\omega_{17} b$$
$$+ \gamma_1 (1 - \gamma_2)\omega_{18} b$$
$$+ \gamma_2 (1 - \gamma_1)(\omega_{19} c + \omega_{20} a + \omega_{21} b)$$
$$+ \gamma_1 \gamma_2 (\omega_{22} a + \omega_{23} b),$$

where $\sum_i (1 - \gamma_1)(1 - \gamma_2)\omega_i + \sum_j \gamma_1 (1 - \gamma_2)\omega_j + \sum_l \gamma_2 (1 - \gamma_1)\omega_l + \sum_m \gamma_1 \gamma_2 \omega_m = 1$,

$\sum_{i=1}^{2} (1 - \gamma_1)(1 - \gamma_2)\omega_i + \gamma_1 (1 - \gamma_2)\omega_3 + \sum_{l=4}^{6} \gamma_2 (1 - \gamma_1)\omega_l + \gamma_1 \gamma_2 \omega_7 = P(G_c)$,

$(1 - \gamma_1)(1 - \gamma_2)\omega_8 + \sum_{j=9}^{10} \gamma_1 (1 - \gamma_2)\omega_j + \sum_{l=11}^{15} \gamma_2 (1 - \gamma_1)\omega_l + \gamma_1 \gamma_2 \omega_{16} = P(G_{dis1})$ and

$(1 - \gamma_1)(1 - \gamma_2)\omega_{17} + \gamma_1 (1 - \gamma_2)\omega_{18} + \sum_{l=19}^{21} \gamma_2 (1 - \gamma_1)\omega_l + \sum_{m=22}^{23} \gamma_1 \gamma_2 \omega_m = P(G_{dis2})$.

Here we define three probabilities (i) $P^{(3)}_{2,AB}$ of observing exactly two coalescences for three lineages in a population $AB$ , (ii) $P^{(3)}_{1,AB}$ of exactly one coalescence for three lineages in a population $AB$ and (iii) $P^{(3)}_{0,AB}$ of no coalescences for three lineages in population $AB$ (Rosenberg 2002). Also, note that $P^{(3)}_{2,AB} + P^{(3)}_{1,AB} + P^{(3)}_{0,AB} = 1$.

$$P^{(3)}_{0,AB} = e^{-3\frac{2}{\theta_{AB}}(h_{ABC} - h_{AB})},$$
$$P^{(3)}_{1,AB} = 1 - \frac{3}{2} e^{-\frac{2}{\theta_{AB}}(h_{ABC} - h_{AB})} + \frac{1}{2} e^{-3\frac{2}{\theta_{AB}}(h_{ABC} - h_{AB})},$$
$$P^{(3)}_{2,AB} = \frac{3}{2} e^{-\frac{2}{\theta_{AB}}(h_{ABC} - h_{AB})} - \frac{3}{2} e^{-3\frac{2}{\theta_{AB}}(h_{ABC} - h_{AB})}.$$

These probabilities that involve coalescence of three branches can be conveniently expressed in terms of $P^{(2)}_{0,\,AB} = e^{-\frac{2}{\theta_{AB}}(h_{ABC} - h_{AB})}$, i.e.

$$P^{(3)}_{0,AB} = (P^{(2)}_{0,\,AB})^3,$$
$$P^{(3)}_{1,AB} = 1 - \frac{3}{2} P^{(2)}_{0,\,AB} + \frac{1}{2} (P^{(2)}_{0,\,AB})^3,$$
$$P^{(3)}_{2,AB} = \frac{3}{2} P^{(2)}_{0,\,AB} - \frac{3}{2} (P^{(2)}_{0,\,AB})^3,$$
$$P^{(2)}_{0,\,B} = e^{-\frac{2}{\theta_B}(h_{AB} - h_i)},$$

$$P^{(2)}_{0,C} = e^{-\frac{2}{\theta_c}(h_{ABC} - h_i)}.$$

Now, we can establish the equality constraints between different model weights:

$$\omega_1 = \omega_{22} = 1 - P^{(2)}_{0,AB},$$
$$\omega_2 = \omega_8 = \omega_{17} = \frac{1}{3}P^{(2)}_{0,AB},$$
$$\omega_3 = \omega_{10} = \omega_{18} = P^{(2)}_{0,C},$$
$$\omega_4 = \omega_{12} = \omega_{19} = \frac{1}{3}P^{(2)}_{0,B} P^{(3)}_{2,AB},$$
$$\omega_5 = \omega_{13} = \omega_{20} = \frac{1}{3}P^{(2)}_{0,B} P^{(3)}_{1,AB},$$
$$\omega_6 = \omega_{15} = \omega_{21} = \frac{1}{3}P^{(2)}_{0,B} P^{(3)}_{0,AB},$$
$$\omega_7 = \omega_{16} = \omega_{23} = \frac{1}{3}P^{(2)}_{0,AB},$$
$$\omega_9 = 1 - P^{(2)}_{0,C},$$
$$\omega_{11} = (1 - P^{(2)}_{0,B})(1 - P^{(2)}_{0,AB}),$$
$$\omega_{14} = (1 - P^{(2)}_{0,B})P^{(2)}_{0,AB}.$$

The only weights that are not constrained include ($\omega_9$, $\omega_{11}$, $\omega_{14}$) and are unique to discordant topology $G_{dis1}$ mixture component, thus:

$$\begin{aligned}\mathbb{E}[T_{MRCA}] = {} & (1 - \gamma_1)(1 - \gamma_2)(\omega_1 a + 3\omega_2 b) \\ & + \gamma_1(1 - \gamma_2)(\omega_9 a + 3\omega_3 b) \\ & + \gamma_2(1 - \gamma_1)(\omega_{11} d + 3\omega_4 c + (3\omega_5 + \omega_{14})a + 3\omega_6 b) \\ & + \gamma_1\gamma_2(\omega_1 a + 3\omega_2 b),\end{aligned}$$

$$\mathbb{E}_{G_c}[T_{MRCA}] = \frac{(1-\gamma_1)(1-\gamma_2)(\omega_1 a + \omega_2 b) + \gamma_1(1-\gamma_2)\omega_3 b + \gamma_2(1-\gamma_1)(\omega_4 c + \omega_5 a + \omega_6 b) + \gamma_1\gamma_2\omega_7 b}{(1-\gamma_1)(1-\gamma_2)(\omega_1 + \omega_2) + \gamma_1(1-\gamma_2)\omega_3 + \gamma_2(1-\gamma_1)(\omega_4 + \omega_5 + \omega_6) + \gamma_1\gamma_2\omega_7},$$

$$\mathbb{E}_{G_{dis1}}[T_{MRCA}] = \frac{(1-\gamma_1)(1-\gamma_2)\omega_2 b + \gamma_1(1-\gamma_2)(\omega_9 a + \omega_3 b) + \gamma_2(1-\gamma_1)(\omega_{11} d + \omega_4 c + \omega_5 a + \omega_{14} a + \omega_6 b) + \gamma_1\gamma_2\omega_7 b}{(1-\gamma_1)(1-\gamma_2)\omega_2 + \gamma_1(1-\gamma_2)(\omega_9 + \omega_3) + \gamma_2(1-\gamma_1)(\omega_{11} + \omega_4 + \omega_5 + \omega_{14} + \omega_6) + \gamma_1\gamma_2\omega_7},$$

$$\mathbb{E}_{G_{dis2}}[T_{MRCA}] = \frac{(1-\gamma_1)(1-\gamma_2)\omega_2 b + \gamma_1(1-\gamma_2)\omega_3 b + \gamma_2(1-\gamma_1)(\omega_4 c + \omega_5 a + \omega_6 b) + \gamma_1\gamma_2(\omega_1 a + \omega_7 b)}{(1-\gamma_1)(1-\gamma_2)\omega_2 + \gamma_1(1-\gamma_2)\omega_3 + \gamma_2(1-\gamma_1)(\omega_4 + \omega_5 + \omega_6) + \gamma_1\gamma_2(\omega_1 + \omega_7)}.$$

Under MSCig model $\mathbb{E}[T_{MRCA}]$ is expressed as follows

$$\mathbb{E}[T_{MRCA}] = (1-\gamma)(\omega_1 a + \omega_2 b + \omega_3 c + \omega_4 c + \omega_5 d) + \gamma\,\omega_6 d$$
$$+ (1-\gamma)(\omega_7 b + \omega_8 c + \omega_9 d) + \gamma\,(\omega_{10} c + \omega_{11} d)$$
$$+ (1-\gamma)(\omega_{12} b + \omega_{13} c + \omega_{14} d) + \gamma\,\omega_{15} d,$$

where $\sum_i (1-\gamma)\omega_i + \sum_j \gamma\omega_j = 1$, $\sum_{i=1}^{5}(1-\gamma)\omega_i + \gamma\omega_6 = P(G_c)$, $\sum_{i=7}^{9}(1-\gamma)\omega_i + \sum_{j=10}^{11}\gamma\omega_j = P(G_{dis_1})$ and $\sum_{i=12}^{14}(1-\gamma)\omega_i + \gamma\omega_{15} = P(G_{dis_2})$ and coalescence probabilities:

$$P^{(2)}_{0,AB} = e^{-\frac{2}{\theta_{AB}}(h_{ABC}-h_{AB})},$$
$$P^{(2)}_{0,ABC} = e^{-\frac{2}{\theta_{ABC}}(h_{ABCX}-h_{ABC})},$$
$$P^{(3)}_{0,ABC} = (P^{(2)}_{0,ABC})^3,$$
$$P^{(3)}_{1,ABC} = 1 - \tfrac{3}{2}P^{(2)}_{0,ABC} + \tfrac{1}{2}(P^{(2)}_{0,ABC})^3,$$
$$P^{(3)}_{2,ABC} = \tfrac{3}{2}P^{(2)}_{0,ABC} - \tfrac{3}{2}(P^{(2)}_{0,ABC})^3,$$

$$\omega_1 = (1 - P^{(2)}_{0,AB})(1 - P^{(2)}_{0,ABC}),$$
$$\omega_2 = \omega_7 = \omega_{12} = \tfrac{1}{3}P^{(2)}_{0,AB}P^{(3)}_{2,ABC},$$
$$\omega_3 = (1 - P^{(2)}_{0,AB})P^{(2)}_{0,ABC},$$
$$\omega_4 = \omega_8 = \omega_{13} = \tfrac{1}{3}P^{(2)}_{0,AB}\,P^{(3)}_{1,ABC},$$
$$\omega_5 = \omega_9 = \omega_{14} = \tfrac{1}{3}P^{(2)}_{0,AB}P^{(3)}_{0,ABC},$$
$$\omega_6 = \omega_{11} = \omega_{15} = \tfrac{1}{3}P^{(3)}_{0,ABC}.$$

Finally, the applying the weight constraints and mixture weight normalization yields:

$$\mathbb{E}[T_{MRCA}] = (1-\gamma)(\omega_1 a + 3\omega_2 b + (\omega_3 + 3\omega_4)c + 3\omega_5 d) + \gamma(\omega_4 c + 3\omega_6 d),$$
$$\mathbb{E}_{G_c}[T_{MRCA}] = \frac{(1-\gamma)(\omega_1 a + \omega_2 b + \omega_3 c + \omega_4 c + \omega_5 d) + \gamma\,\omega_6 d}{(1-\gamma)(\omega_1 + \omega_2 + \omega_3 + \omega_4 + \omega_5) + \gamma\,\omega_6},$$
$$\mathbb{E}_{G_{dis1}}[T_{MRCA}] = \frac{(1-\gamma)(\omega_2 b + \omega_4 c + \omega_5 d) + \gamma(\omega_{10} c + \omega_6 d)}{(1-\gamma)(\omega_2 + \omega_4 + \omega_5) + \gamma(\omega_{10} + \omega_6)},$$
$$\mathbb{E}_{G_{dis2}}[T_{MRCA}] = \frac{(1-\gamma)(\omega_2 b + \omega_4 c + \omega_5 d) + \gamma\omega_6 d}{(1-\gamma)(\omega_2 + \omega_4 + \omega_5) + \gamma\omega_6}.$$

### Supplementary tables

| Supplementary Table One: Input Used to Generate Extended Newick Trees with the msci-create function of BPP under Each Demographic Scenario | |
|---|---|
| Demographic scenario | Input |
| Ghost Introgression from G to B | `tree (G,(C,(B,A)T)R)Z;`<br>`hybridization Z G, T B as S H tau=no, yes phi=0.30` |
| Inflow Introgression from C to A | `tree (C,(B,A)T)R;`<br>`hybridization R C, T A as S H tau=no, yes phi=0.30` |
| Outflow Introgression from A to C | `tree (C,(B,A)T)R;`<br>`hybridization T A, R C as S H tau=no, yes phi=0.30` |
| *The species tree (C,(A,B)) was provided without alteration to test the “no introgression” model. | |
